# Quantitative analysis of three-dimensional cell organisation and concentration profiles within curved epithelial tissues

**DOI:** 10.1101/2022.05.16.492131

**Authors:** Chaitra Prabhakara, Krishnan Swaminathan Iyer, Madan Rao, Timothy E Saunders, Satyajit Mayor

## Abstract

Organogenesis involves folding of flat epithelial tissues into three-dimensional (3D) shapes. Quantitative analysis in 3D of the concentration gradients of biochemical and mechano-chemical cues that shape the tissue has remained a challenge, due to the complex tissue geometries. Towards addressing this, we present a methodology that transforms the cartesian images acquired by high-resolution confocal microscopy of curved tissues into a laminar organisation from the outer-most tissue surface. We further detail a data-based intensity correction method to account for the intensity variations that arise as a consequence of the global geometry. Applying our approach to the dome-shaped *Drosophila* wing disc, we quantitatively estimate the concentration profiles of different biochemical signals, including the Wingless morphogen gradient. The laminar data-organisation method also enabled visualisation of apical and basal layers facilitating an accurate 3D reconstruction of cell shapes within the pseudostratified epithelium. We find that the columnar disc proper cells have irregular 3D shapes and undergo frequent apico-basal cell intercalations, concomitantly leading to cell neighbour exchanges along the apico-basal axis. The workflow of image processing described here may be employed to quantify 3D concentration gradients and 3D cellular organisation in any layered curved tissue, provided a marker describing the reference surface manifold is available.

## Introduction

Epithelial tissues display a large diversity in both function and form as columnar, cuboidal, or squamous shapes that are organised in mono or multilayers. Typical epithelial cell cultures consist of a monolayer of cells which grow and divide, with cell dynamics typically restricted to the two-dimensions (2D) of the cell culture dish. However, *in vivo*, most epithelia are not just flat sheets of cells and can be considerably curved with radius of curvature comparable to a few cell lengths (reviewed in (Gómez-Gálvez *et al*., 2021)). Like an origami crane emerging from a flat sheet of paper with simple folding rules, complex three-dimensional (3D) epithelial structures observed during development are formed using different biochemical and mechanical cues (reviewed in (Hamant and Saunders, 2020)). Morphogenetic processes driven by cell shape changes, cell rearrangements and oriented cell divisions lead to out-of-plane bending of epithelia into folds, ridges and tubes (reviewed in (Guillot and Lecuit, 2013; Lemke and Nelson, 2021)). Local mechanical properties and their effects on tissue topology can also influence downstream cell fate decisions by shaping morphogen fields (Shyer *et al*., 2013; Manning *et al*., 2015). Studying the native epithelia in 3D can provide insights into how cells are organised and allow quantitative measurement of the biochemical concentration profiles that guide organogenesis.

Dramatic improvements in microscopy over the last 20 years have resulted in the resurgence of the field of developmental biology (reviewed in (Keller, 2013)). Modern microscopy techniques aid in multi-channel, multi-timepoint imaging to achieve single cell resolution in a wide field of view that covers most of the tissue of interest (Krzic *et al*., 2012). Such techniques have allowed visualisation of, for example, (i) morphogen gradients (Gregor *et al*., 2007; Kicheva *et al*., 2007; Durrieu *et al*., 2018), (ii) cell dynamics leading to tissue formation or repair (Rauzi *et al*., 2015; Dye *et al*., 2017; Park *et al*., 2017), (iii) selective localisation of PCP pathway components (Devenport and Fuchs, 2008; Aigouy *et al*., 2010), and (iv) the underlying actin-myosin meshwork bringing about epithelial sheet rearrangements (Martin, Kaschube and Wieschaus, 2009; Munjal *et al*., 2015; Streichan *et al*., 2018). Most of these processes have been quantitatively studied using maximum intensity projections (MIP) of the curved tissue onto a 2D plane. Simple MIP has a fundamental drawback: noise, especially from within cells, accumulates in the projected image and thus, compromises the signal-to-noise ratio. To address this limitation, tools have been developed to project only the tissue layer from where the intensity arises, such as StackFocuser (Umorin, 2006), SurfCut (Erguvan *et al*., 2019), MinCostZSurface, Premosa (Blasse *et al*., 2017) and Extended Depth of Field (Forster *et al*., 2004). These approaches work best when studying processes near or at the apical surface. However, limitations to such 2D projections are observed when quantifying signals from the entire apico-basal length of the cell. Despite the rapid advancement of 3D imaging technologies, these methods are still not commonly used to process and quantify such data.

Epithelial cells interact with their neighbours through tight junctions at the apical side and bind to basal lamina and extracellular matrix on the basal side (reviewed in (Rodriguez-Boulan and Macara, 2014)). The arrangement of epithelial cells from an apical perspective has been extensively studied (Baena-López, Baonza and García-Bellido, 2005; Classen *et al*., 2005; Gibson *et al*., 2006; Aigouy *et al*., 2010). Stereotypical changes in apical cell shape promotes tissue remodelling during invagination and extension. Apical constriction, as observed in *Drosophila* embryonic mesoderm invagination and neurulation in vertebrates, is conceptualised as the conversion of taller columnar cells to shorter frusta or bottle-like cells that have a smaller apical area and a larger basal area (reviewed in (Lecuit and Lenne, 2007)). Convergent extension brought about by cell intercalations, wherein a group of four cells alter their neighbours through T1 transitions, is observed during *Drosophila* germband extension (Bertet, Sulak and Lecuit, 2004) and tube extension of *Drosophila* tracheal system (Ribeiro, Neumann and Affolter, 2004).

Is the cell organisation observed on the apical side a representation of how cells are arranged in 3D? Though more difficult to image and analyse, recent studies (Rupprecht *et al*., 2017; Sun *et al*., 2017; Gómez-Gálvez *et al*., 2018; Gómez *et al*., 2021) have shown that cell intercalations occur not only along the apical surface in time but along the apico-basal axis in space. Such an organisation, unlike apical constriction, results in different neighbours along the apico-basal axis. Depending on the anisotropy of apical and basal cell surfaces of the curved epithelium, the cells can be arranged as prisms or frusta or a combination of frusta and scutoids (Gómez-Gálvez *et al*., 2018). Understanding the nature of the 3D cell packing and alterations in cell neighbour changes is important, as these physical properties can affect the packing density, cell-cell transport of morphogens (Müller *et al*., 2013) and cell-cell signalling (Yaron, Cordova and Sprinzak, 2014; Corson *et al*., 2017), and thereby potentially play a role in altering cell fate determinations.

Here, we develop a framework to organise confocal imaging data of curved epithelia as layers with respect to the outermost surface of interest. We use the curved pseudostratified epithelium of the wing imaginal disc as a model system. We describe a method to correct the intensity bias brought about by imaging artefacts due to sample geometry and use the corrected intensity to quantify intensity distribution of biochemical signals, such as morphogen gradients, in 3D within each apico-basal layer with respect to distance from a reference plane. Such a method of organisation also helps in separating the apical and basal surfaces, reconstructing 3D cellular organisation and quantifying morphometric features such as cell area and cell neighbours as a function of apico-basal axis.

## Results

### Organisation of cells in the curved epithelium of the wing imaginal disc

The wing disc is a folded epithelial sac with two kinds of cells: the outer squamous epithelial cells called the peripodial cells and a single layer of columnar cells called the disc proper (Figure 1A). We focus on the wing pouch of the wing disc (green region in Figure 1A), which is an elliptical dome with a dense mesh of columnar cells. Third instar (96-110 h AEL) larval wing discs were dissected and mounted with spacers to image the tissue in 3D using confocal microscopy (Methods). Images of wing disc expressing different markers were acquired slice-by-slice to obtain 40-50μm of optical sectioning, starting from the top of the dome to the bottom (Schematic in Figure 1B, Figure S1A). Imaging wing discs labelled with an apical adherens junction marker, *E-Cadherin* (Tepass *et al*., 1996), describes the manifold of cell arrangements within the curved tissue (Figure S1). Given the dome shaped nature of the epithelia, the top Z slices mostly show the apical parts of the cells in the centre, while the remaining imaging planes display the apical cells towards the tissue periphery (Figure S1A). Orthogonal views of apically localised *E-Cadherin* describe the shape of the wing disc to be a hemi-ellipsoid with curvatures seen along both major and minor axes (Figure S1B).

**Figure 1:**
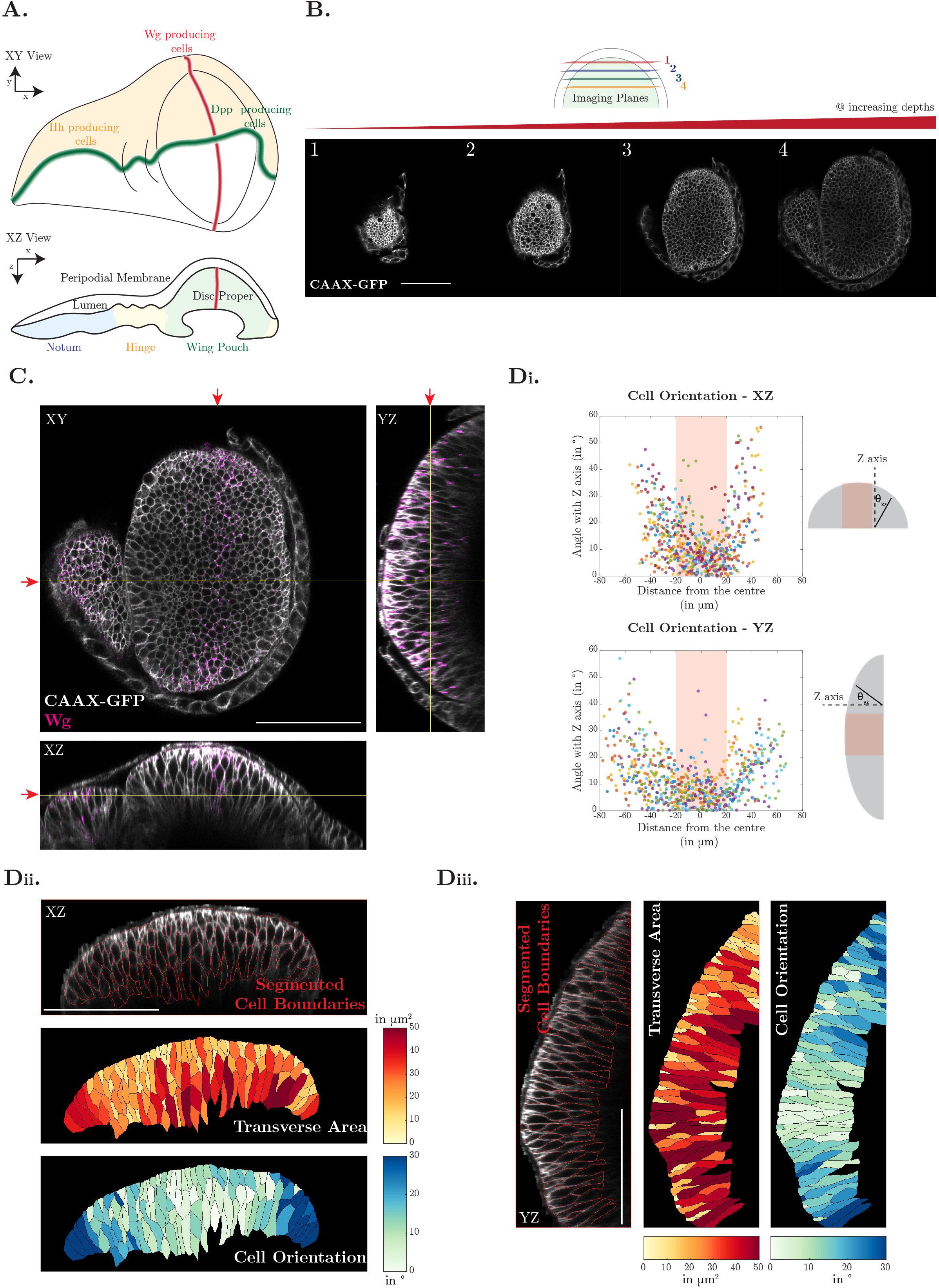
Orientation of cells within the curved epithelium of wing disc. A: Schematic representation of the three-dimensional wing imaginal disc. Y axis (major axis) represents the *Wg* producing cells, X axis (minor axis) represents cells at increasing distance from the producing cells, Z axis represents the depth of the cells. Expression domains of key signalling molecules (*Wg*, *Dpp* and *Hh*) and fate map (Notum, Hinge, Wing Blade) are also indicated. B: Z sections of *CAAX-GFP* expressing wing imaginal disc. The different Z sections (1-4) show that each plane has information about cells at different heights. For example, the rim of (4) has cells with smaller apical area, while the central region shows cells with larger medial/basal area of the pseudostratified epithelium. C: Orthogonal views of wing disc labelled with *CAAX-GFP* (cyan) and *Wg* (magenta) shows curvature along both minor and major axis of the wing disc. The red arrow marks the section represented in XY, YZ, XZ projection. D: Segmentation of transverse section of wing disc XZ (ii) and YZ (iii) helps in estimating the angle of orientation of cells (i). The heat map in ii and iii show that cells along the periphery are more tilted with respect to the vertical illumination axis (Z) compared to the cells in the centre. The cell orientation quantification shown in (i) describes that the cells are tilted away from the centre (shaded orange) across both axes. Cells are more tilted away from the centre across the minor axis (XZ) than the major axis (YZ). Colours in the plot in (i) represent different transverse slices. Number of cells: 628 cells from 7 transverse slices (XZ) and 690 cells from 6 transverse slices (YZ) from 2 samples. On an average, the 2D cross-sectional orientation of cells in XZ and YZ is 14.7±12.2° and 12.6±10.1° (mean±sd) respectively. Scale bar: 50μm

A simple approach to visualise and analyse 3D datasets is to use a 2D planar projection in the direction of the imaging axis, defined by the imaging platform as the axis along which the sample is optically sectioned. Although dimensionality reduction is useful in reducing the data size and enabling additional analysis routines like cell segmentation and cell tracking, which currently are more robust in 2D compared to 3D, much of the spatial information is lost (Heemskerk and Streichan, 2015). As apical markers label only a part of the cell, the intensity projection (Umorin, 2006) of apical markers onto a 2D plane (for a tissue with a single layer of epithelial cells) is feasible and informative. Cell segmentation conducted on such projected images (Figure S1C(i), Methods) revealed the average apical area of wing disc columnar cells to be 4.4±2.0 μm^2^ (Figure S1D). Additionally, spatial information can be retrieved from the depth map (or *z-depth* map) of maximum intensity of *E-Cadherin* that describes the contour of the reference apical surface: a smooth and continuous increase in *z-depth* is seen as we move away from the centre (Figure S1C(ii), Methods). Thus, the apical marker describes the outermost surface of cells embedded within the Euclidean 3D space.

The approach to studying 3D datasets becomes more complex when studying markers that label the entire apico-basal length of cells in a curved tissue. To analyse the 3D architecture of the epithelial cells, wing discs expressing the plasma membrane marker CAAX-GFP were evaluated (Figure 1B, Movie S1). On slicing through the sample from the top of the dome to the bottom, it is evident that the cell organisation is more complex than the apical surface alone. Orthogonal views of CAAX-GFP-labelled wing disc clearly show the tissue curvature along both the minor and major axes (Figure 1C). From these views, we also see that: (i) CAAX-GFP labels the entire length of the cells; (ii) cells are elongated along the vertical axis; (iii) cells are densely packed; and (iv) the tissue is pseudostratified with multiple cross-sectional cell shapes (Movies S2-S3). The 2D transverse section of flat or curved tissue with columnar epithelial cells usually has a single cell along any axis perpendicular to the surface. In this case of pseudostratified cells, multiple cell boundaries were encountered while moving along any axis perpendicular to the surface of each 2D transverse section (Figure 1D). This is unlike the orderly packed columnar epithelial cells previously observed in the early *Drosophila* embryo (Rupprecht *et al*., 2017).

To study the cellular organisation of this pseudostratified epithelium, we quantified the area and orientation of cells in each axis view. The angle of cell orientation, defined as the angle between the long axis of the segmented cell with the vertical (Z) axis, is overlaid as a heatmap on the segmented cells (Figure 1D(ii), 1D(iii)). The cell orientation angles show that while the cells in the centre of the disc have lower orientation angles, cells in the periphery have larger deviations in their angles (Figure 1D(i)). The cells in the periphery are more tilted along the minor axis than along the major axis (Figure 1D(i)). While the transverse section measurements show a trend in cell position and cell orientation, the measurements have a limitation in that the cross-sectional view encompasses only a part of the entire cell and not the whole cell due to the curved pseudostratified nature of the epithelium. Cells within the pseudostratified disc proper tissue appear to be aligned perpendicular to the outermost surface.

### Transforming 3D image datasets into laminar organisation from the outermost surface

Simple 2D projections, like that for *E-Cadherin*, fail for CAAX-GFP or markers which label cells along the length of the apico-basal axis as such projections cannot describe clear cell boundaries, especially for a curved pseudostratified epithelium. It is therefore useful to transform the optical slices obtained along the axes defined by the imaging platform during data acquisition, into layers defined by the perpendicular distance from the outermost surface. In this section, we discuss the methodology used to perform this transformation with the steps outlined in Figure 2A and detailed in Methods.

**Figure 2:**
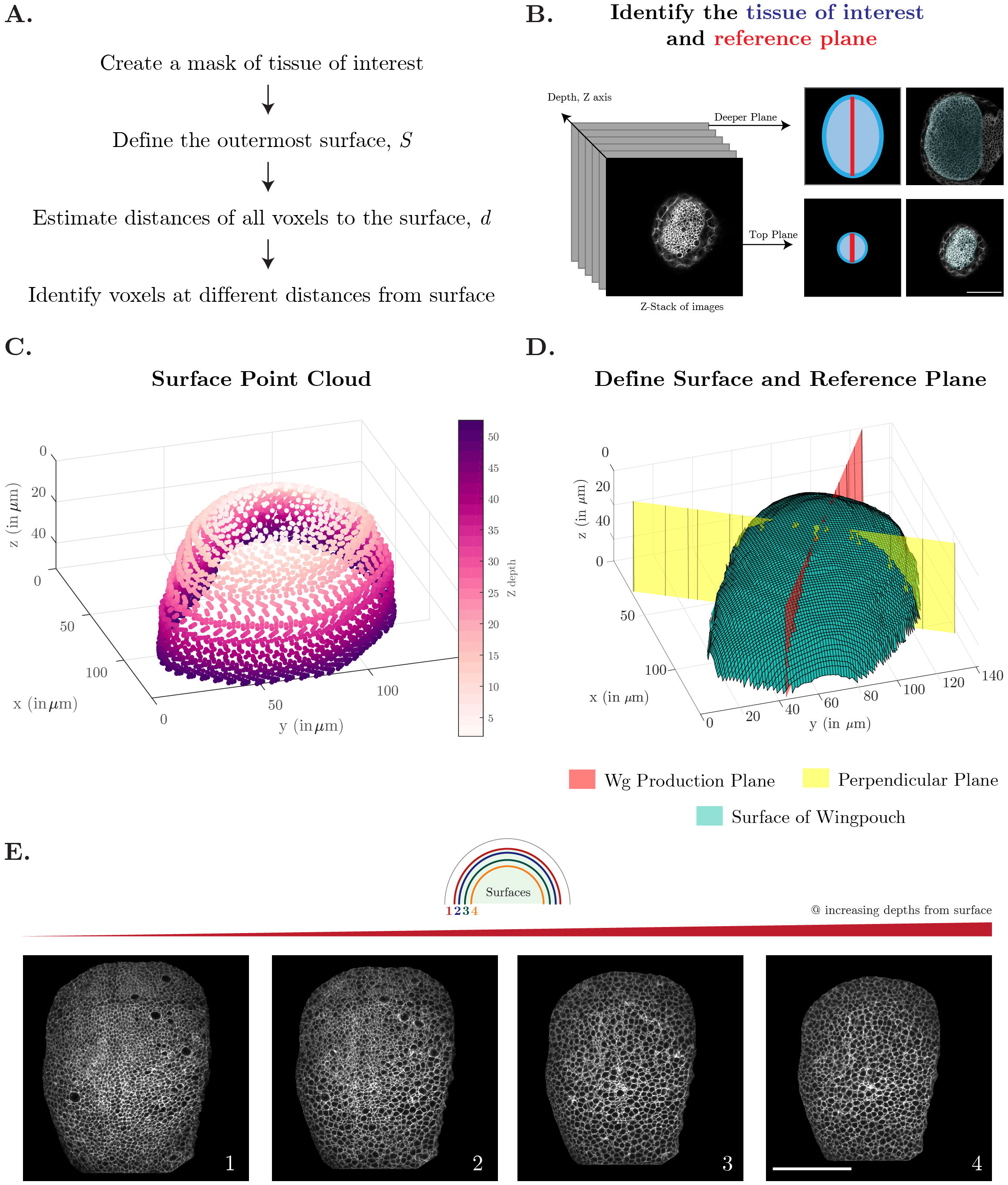
Workflow to reorganise 3D image data as layers with respect to the outermost surface. A-E: The workflow to restructure 3D image data into laminar organisation with respect to the outermost surface is summarised in A and output is shown in E. The binary mask of the tissue of interest (blue overlay in B) is smoothened and interpolated to define a point cloud describing the outermost surface of wing disc (C, D). The *Wg* production plane (red) and the perpendicular plane (yellow) is defined for each wing disc. For any evaluation voxel inside the 3D image stack, the shortest distance from the surface (*d*) was computed to organise the 3D image data as layers with respect to the outermost surface. E: Projection of each layer of CAAX-GFP labelled wing disc moving inwards from the outermost surface as described in the schematic. Each layer is 0.5μm thick. See Methods for more information. Scale Bar: 50 μm.

In the process of confocal imaging, along with the tissue of interest (*i.e*., disc proper cells), other tissues in the proximity, such as peripodial membrane and hinge cells, are inevitably captured. Peripodial cells are not at a constant distance away from the disc proper cells (see Figure S1B). Therefore, a simple Z shift is not sufficient to remove the peripodial layer of cells. To separate the tissue of interest from other tissues, masks were hand-drawn, and the outlines of the masks were smoothened (Figure 2B). Subsequently, the outlines of the masks were linearly interpolated to extract the apical-most surface, *S* (Figure 2C, 2D). This point cloud *S* defines a curved 3D region. The distance of all points within the wing disc to the outermost surface, *S*, was computed next. The surface, *S*, was approximated to be a series of small line segments (piecewise linear chordal segments) (D’Errico, 2021). The Euclidean distance of each point P within the wing disc to these piecewise line segments was calculated. The minimum of this set was estimated as the distance *D_pp’_*, with *P*’ being the nearest point on *S,* for every P (Methods, Figure S2A). Using this method, each pixel within the tissue of interest could be categorised by its distance from the outermost surface.

Each isosurface at different distances from the outermost surface can be analysed to estimate intensity profiles of various fluorescently labelled markers or can be projected onto a 2D surface for visualisation of cell organisation along the apico-basal axis (Methods). An example of the projection of layers onto a 2D surface is shown in Figure 2E and Movie S4. This 3D geometrical categorisation overcomes the problems of simple 2D planar projection observed for a curved pseudostratified epithelium. Such a transformation improves the visualisation of apical surfaces as distinct from basal surfaces. Unlike the confocal stacks, cells in the centre or periphery in the transformed planes are now at the same height. The framework outlined above provides a clear method for comparing cells at similar heights within the tissue, rather than by their distance to the (arbitrary) imaging plane. Using this methodology, any curved sample can be deconvolved into layers, equidistant from the outermost surface.

### Parameterising the dependence of sample geometry on fluorescence intensity

Depth of imaging is limited in confocal microscopy (Jonkman *et al*., 2020). A peculiarity which stands out in 3D datasets of curved samples imaged using confocal microscopy is the intensity gradient observed within each layer. Pseudo-coloured images of each layer of wing discs labelled with CAAX-GFP show an intensity gradient where the centre of the sample is brighter, with the intensity falling off gradually as we move away from this centre (Figure 3A). However, as CAAX-GFP is driven by a Ubiquitin promoter, the expectation is that all cells will have similar expression levels of CAAX-GFP. Moreover, on transforming the imaging data as layers from the outermost surface, each layer has information from cells at the same depth along the apico-basal axis. Therefore, each surface layer is expected to have similar distributions of intensities and the observed intensity variations are likely linked to the curvature-based artefacts.

**Figure 3:**
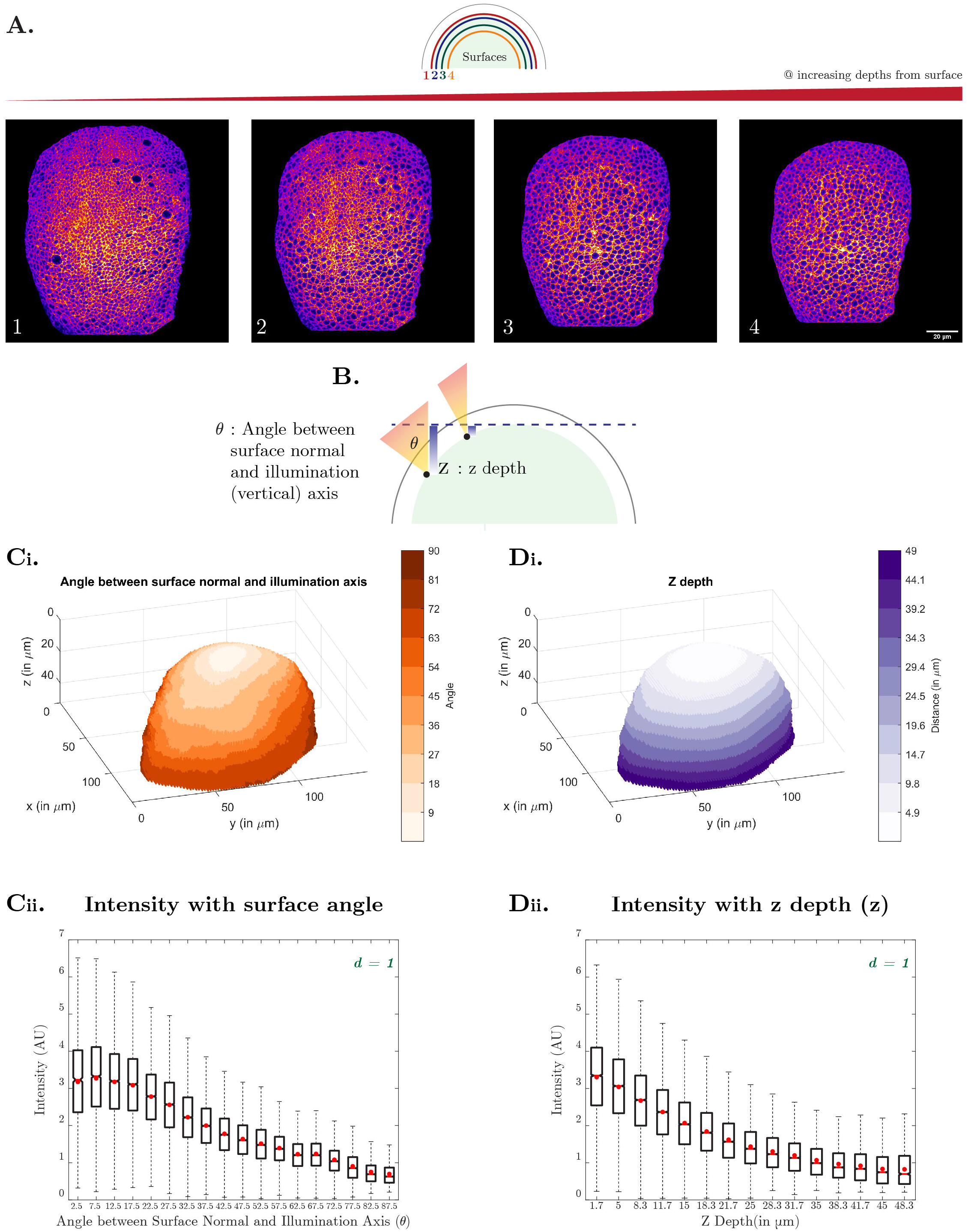
Imaging and sample geometry parameters affecting intensity within the curved epithelium of wing disc. A: Pseudo-coloured projection images of each layer of CAAX-GFP labelled wing disc moving inwards from the outermost surface as described in the schematic. Each layer is 0.5μm thick. See Methods for more information. This figure is related to Movie S4 and Fig 2E. Scale Bar: 20 μm. B: Schematic describing sample geometry parameters: imaging depth (*z depth*) and surface angle (*θ*) C-D: Normalised intensity (Intensity/Mean Intensity) expressing CAAX-GFP shows that intensity reduces with increasing surface angle (Ci, Cii) as well as increasing imaging depth (Di, Dii). The boxplot refers to intensity distributions from voxels identified in the topmost layer (*d* = 1 is the apical most layer of 4.8μm thickness) of four wing discs.

What factors could contribute to an intensity gradient? In confocal microscopy, the signal levels are profoundly sensitive to scattering of excitation and emission light. We see that across wing discs, the intensity of CAAX-GFP reduces with sample depth (distance from the outermost surface, Figure S3A(i)). While scattering of light affects the recorded intensity, it alone cannot explain the variations in intensities observed within each layer (Figure 3A, S3Aii). The heterogeneity of intensities as measured by coefficient of variation of intensities is larger in the apical layers and reduces with increased sample depth (Figure S3A(ii)). To interrogate the sources of intensity variations within layers, we next measured the dependence of intensity on two factors related to sample geometry: imaging depth (*z-depth*) and the surface angle (*θ*) (Figure 3B). Imaging depth is defined as the vertical distance from the first imaging plane. Surface angle is defined as the included angle between the surface normal and the vertical illumination (Z) axis (Methods). Note that the surface angle is not the same as polar angle. The larger the angle *θ*, the more oblique is the surface with respect to the illumination axis. Similar to a simulated hemi-ellipsoid (Figure S3B), points closer to the centre of the wing disc have small deviations in surface angles with respect to the illumination axis, while those in the periphery show larger deviations (Figure 3Ci, S3C). The CAAX-GFP intensity measured across wing discs reduces as the surface angle increases within each surface layer (Figure 3Cii). As expected for a hemi-ellipsoid (Figure S3B), points closer to the centre of the wing disc have lower *z-depth* values while those in the periphery show larger *z-depths* (Figure 3Di, S3C). A reduction in CAAX-GFP intensity was also observed with increasing imaging depth for each surface layer (Figure 3Dii). These observations indicate that scattering in combination with the curved sample architecture contribute to the observed intensity variations. The range of *z-depths* spanning the outermost apical layer is maximum and this range reduces in basal layers. This explains why the variation in observed intensities is highest in the apical layers and lower in deeper layers (Figure S3Aii).

Within curved tissues, if cells are aligned according to the curvature (Figure 1D), then the orientation of cells with respect to the imaging axis contributes to the curvature related artefacts in the recorded intensities. We provide a simple explanation for this in Supplementary Note 1. Consider a curved tissue with fluorescently labelled apical surface with surface angle *θ* (Figure S4Ai). The average intensity of signal recorded from the curved tissue depends on the surface angle, *θ* and the distance of the curved tissue from the recording plane, *z_0_* (Equation 1, Figure S4Aii). We observe that the variation with surface angle is more pronounced if this curved tissue lies at a shorter distance from the recording plane. At larger distances, the contribution of reduction of intensity due to *z-depth* will be larger than the effect of cell orientation. In the above example, we considered a special case where only the apical surface of the cells is fluorescently labelled. However different biochemical reporters are localised to different organelles within the cell volumes (Figure 5). To extend our understanding of the contribution of cell orientation in altering recorded intensities, we considered two extreme scenarios: one in which the molecules are localised to a thin narrow region within the cell (like the apical adherens junction markers in Figure S4B(i)); and another in which the molecules are spread across the volume of the cell (like the nucleus in Figure S4C(i)). On convolving these objects using a point spread function (Supplementary Note 1), we estimated the recorded intensity for each scenario (Figure S4B(ii), S4C(ii)). The effect of orientation on measured intensity matters more for markers localised to thin volumes compared to those distributed in broader volumes (Figure S4B(iii), S4C(iii)). In case of samples with dispersed marker localisations, with thickness greater than the z-resolution, the loss in intensity for any sample orientation is predominantly because of the *z-depth*. For narrower localisations, with thickness less than the z-resolution, both sample orientation and *z-depth* contribute to the recorded intensities. Our empirical observations combined with these theoretical considerations make clear that sample geometry with respect to the imaging axis causes the recorded intensity differences. While these issues have long been known, such considerations are not fully accounted for in most analyses of curved epithelia. The parameters defined above can be used to correct for such observed intensity differences, as we detail next.

### Accounting for the intensity changes due to curvature

To correct the intensity differences, we constructed a data-based normalisation matrix using CAAX-GFP intensities as a reference. This is valid with the assumption that the differences observed in CAAX-GFP intensity are primarily due to the dependence of intensity on the above stated parameters (depth and orientation). Such a correction matrix allows for the comparison of any concentration profile to that of CAAX-GFP gradient and decouples the variation in fluorescence intensity from systematic variations observed due to scattering and sample geometry.

Different normalisation matrices were generated based on the following parameters: distance from the surface (*d*), *z-depth* (*z*) and/or surface angle (*θ*). We classified the normalisation matrices as either: depth-based normalisation matrix using (*d*) and (*z*) or curvature-based normalisation using (*d*) and (*θ*) or 3D normalisation matrix using (*d, z, θ*) as parameters. These matrices were constructed using intensities from multiple wing discs labelled with CAAX-GFP (Methods, Figure 4A, S5A, S5B). An example of intensity distribution from multiple wing discs for the apical layer (*d = 1*) is shown in Figure S5C. The mean of intensities belonging to each (*d*, *z*, *θ*) was used to construct the normalisation look-up-table (Figure 4A). Most of the *z* and *θ* bins were populated by almost all wing disc samples, but we note that some *z* and *θ* bins were only populated by a few (owing to the surface geometry differences across samples). Overall, the normalisation matrix encompasses the effect of intensity changes for a wide range of orientations of the hemi-ellipsoidal sample.

**Figure 4:**
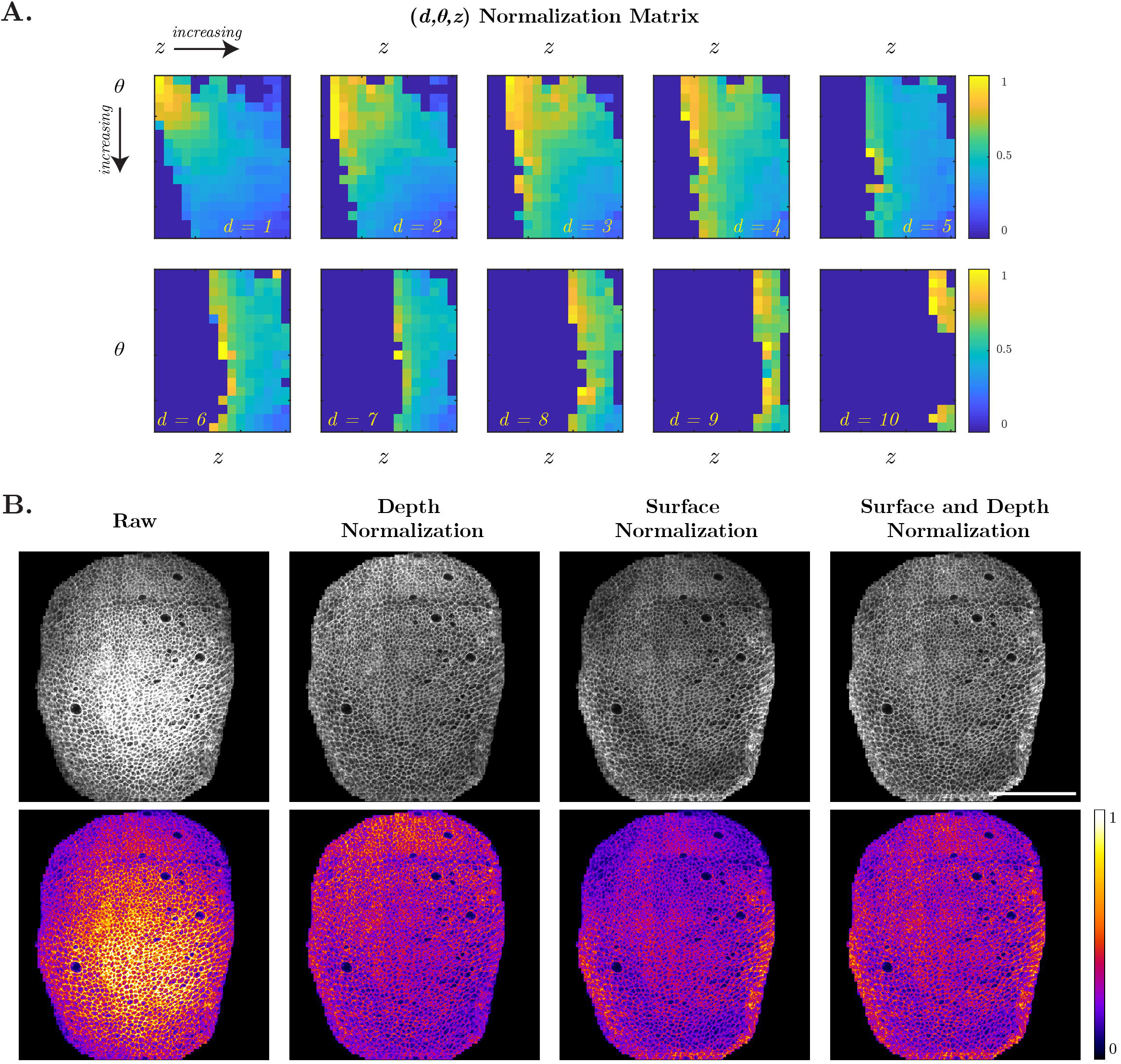
Data based method to correct the intensity changes within curved epithelium. A: 3 parameter normalisation matrix (*d, Z, θ*) constructed from six wing discs expressing CAAX-GFP to remove the intensity bias brought about due to surface geometry B: Intensity bias was corrected using imaging depth alone or surface angle alone or using both imaging depth and sample geometry normalisation. The normalised pseudo-coloured images allow comparison of the intensity distribution of a layer of wing disc (*d* = 1 is the apical most layer of 4.8μm thickness) using different normalisation matrices. Three parameter normalisation works best in equalizing intensity (See more information in Methods). Scale Bar: 50 μm.

For intensity correction, voxels within each wing disc defined by (*d, z, θ*) were corrected using a corresponding factor from the normalisation matrix. While both the 2D normalisations worked well in equalising the intensities across the entire disc, small deviations in intensities were seen with both depth-based and curvature-based normalisation (Figure 4B). A 3D normalisation matrix that included (*d, z, θ*) as parameters functioned the best in correcting the heterogeneities in the signal (Figure 4B, Movie S5). Unlike the symmetric hemi-ellipsoid, the *z-depths* and the surface angles are not symmetric for a wing disc (Figure S3B, S3C). Although closely related, *z* and *θ* together better described the surface of a wing disc (Figure S3C). This approach is limited in correcting the intensity along the extreme edges of the minor axis. This is likely due to the sharper curvature changes observed across the minor axis, more so than along the major axis. Fewer voxels are observed in the highest curvature areas, increasing the probability of having extreme values due to smaller sample size. Therefore, in all the quantifications below the sample extremities have been ignored. Using such a data-based normalisation methodology, the intensity differences brought about due to sample geometry can be accounted for within the limits mentioned above.

### Measuring 3D concentration profiles from the reference plane

Concentration profiles of biomolecules within an epithelium are usually defined from a reference plane. Here, we defined the *Wg* morphogen producing cells as a reference plane. *Wg* is produced by a stripe of cells at the dorso-ventral boundary of the wing disc. The production domain was approximated to be the best-fit plane, *Q,* passing through the points of high intensity in each slice (red plane in Figure 2D) and the perpendicular plane, *R* (yellow plane in Figure 2D), is defined with respect to *Q*. Distances *D_PQ_* and *D_PR_*, the shortest distance of each point, *P*, within the wing disc to the production plane, *Q*, and perpendicular plane, *R*, was measured (Methods). Figure S2B illustrates points within the wing disc divided into several bins based on the distance from the production plane or distance from the perpendicular plane. Estimating concentration profiles across the production plane using *D_PQ_* bins demonstrates how concentration changes as a function of distance from producing cells. Comparing the concentration profiles within *D_PR_* bins provides some understanding on the variability of concentration gradients within each wing disc.

Towards quantifying the intensity as a function of distance from producing cells, each disc was first divided into five layers using distance from the outermost surface (*D_pp’_*). Subsequently, each layer was further divided into thirty bins as a function of distance from producing cells (*D_PQ_*). Mean intensities within each bin were calculated (Methods). All intensities were normalised by the intensity of in the apical-most bin closest to the production plane to facilitate comparison across multiple samples. Figure 5A(iii) shows a representative output of normalised intensity of CAAX-GFP as a function of distance from producing cells and along apico-basal axis. Without correction, we observe a decrease in intensity of CAAX-GFP as a function of distance from producing cells (Figure 5A(iii)). Corrected intensity shows a uniform profile of CAAX-GFP, confirming that the correction code works (Figure 5A(iv)). Correction also helps in reducing the intensity variability observed within each apico-basal layer, allowing an understanding of the actual variation in concentration profiles (Compare Figure S3A(ii) and S5D).

**Figure 5:**
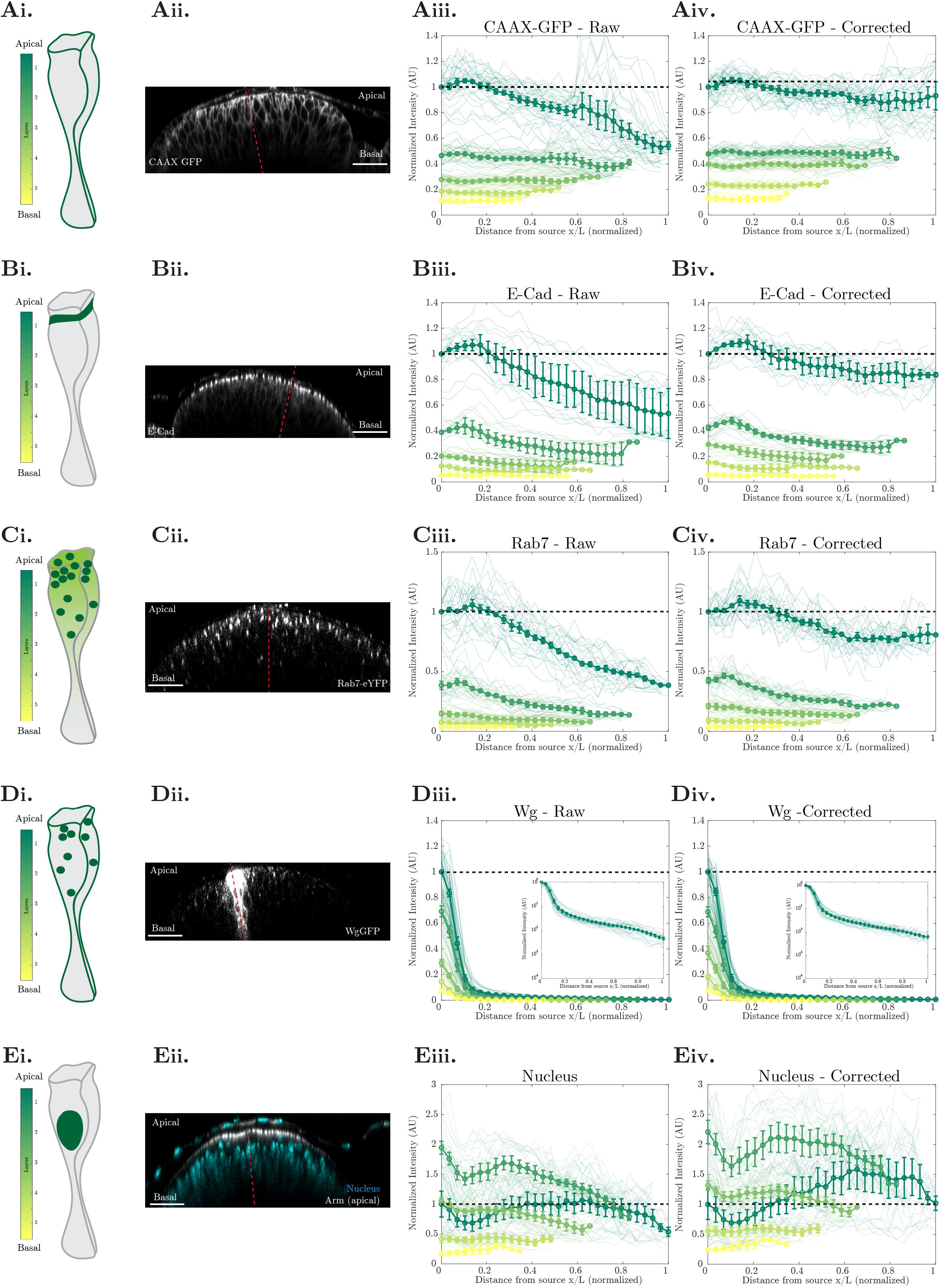
Measuring concentration profiles with respect to a reference plane, Wingless production plane. A-E: Concentration profiles of different probes (A: *CAAX-GFP,* B: *E-cadherin*, C: *Rab7*, D: *Wg-GFP*, E: Nucleus) were measured as a function of distance from producing cells. Cartoon representation of localisation is shown in (i), XZ orthogonal projection in (ii, dotted red line indicates Wg producing cells), raw normalised intensity profiles in (iii) and corrected normalised intensity profiles in (iv). Colours represent the layer number (refer to the LUT depicted in i), darker lines represent the mean and SEM, lighter lines represent individual traces across different stripes from discs (N = 4, 2, 3, 5, 4). Scale bar: 20 μm.

Our 3D analysis was next applied to other samples, such as wing discs labelled with *E-Cadherin, Rab7* and *Wg*. While the uncorrected layers of an example disc expressing *E-Cadherin* shows heterogenous intensities, with the centre being brighter than the periphery, the corrected layers show a near-homogenous distribution of *E-Cadherin* signal (Figure S6A). *E-Cadherin* is slightly elevated next to *Wg* producing cells. While the raw plot shows a gradual decay of *E-Cadherin* intensity as a function of distance from producing cells, in the corrected wing discs the intensity of *E-Cadherin* expression remains nearly constant. This indicates that the decay observed in unprocessed imaging data is primarily due to intensity artefacts and that *E-Cadherin* levels do not decrease away from Wg producing cells (Figure 5B).

Rab GTPases along with their effectors, coordinate vesicle trafficking functions such as formation of vesicles, fusion of vesicles, directing vesicles to target membranes (reviewed in (Caviglia *et al*., 2019)). In wing discs, many Rab proteins are apically enriched. YFP protein trap lines of Rabs showed that those involved in secretion (*Rabs 1, 2, 6, 8*) and those involved in endocytosis (*Rabs 4, 5, 7, 11*) are targeted to apical hub (Dunst *et al*., 2015). Gaining accurate profiles of these endocytic Rab proteins is important as they have consequence in establishing and maintaining morphogen gradients (Wg, Dpp, reviewed in (Gonzalez-Gaitan and Julicher, 2014)). Here, we elucidate the 3D concentration profile of *Rab7-eYFP* (Figure S6B(i)). *Rab7-eYFP* shows apical enrichment (Figure 5C(i), 5C(ii)). While the uncorrected *Rab7* concentration profile shows a gradual decay from the producing cells (Figure 5C(iii)), the intensity corrected *Rab7* shows little gradient across the sample (Figure 5C(iv)). Once again, the differences observed in the uncorrected samples is primarily due to intensity artefacts.

The morphogen *Wg* forms a gradient across the dorso-ventral axis of the wing disc (Neumann and Cohen, 1997). Endogenously tagged *Wg-GFP* (Port *et al*., 2014) shows that *Wg* is enriched in the producing cells (Figure 5D(i), 5D(ii)). Separating *Wg-GFP* into layers shows that *Wg-GFP* is enriched at the producing cells across all layers along the whole apico-basal axis (Figure S6B(ii)). Measuring the average profile of *Wg-GFP* demonstrates that both uncorrected and corrected intensities show a steep decaying curve from the producing cells (Figure 5D(iii), 5D(iv)), consistent with previous overexpression studies (Kicheva *et al*., 2007).

We next evaluated the nuclei profiles within the wing disc. The nucleus is localised in more basal layers (Figure 5E(i), 5E(ii), Movie S6). This is also detected in the quantification, wherein the intensity (of DAPI) in the second layer is more than the intensity recorded in the apical-most layer. We observe that nuclei in cells next to the *Wg* producing cells are localised further away from the apical-most surface (Figure S6B(iii)). The quantification also reflects a dip in intensity right next to the *Wg* producing cells in the apical two layers (Figure 5E(iii)). The corrected intensity for nuclear signal corrects the tissue geometry related decay in intensity observed in cells away from the producing cells (Figure 5E(iv)).

The examples discussed above indicate the necessity of applying intensity correction for an accurate quantification of markers. As the apical layers show the highest variation in intensity in unprocessed images, intensity correction is especially required for quantification of markers localised to apical layers. Additionally, the effect of correction is more prominent for markers which are localised to cells in the centre as well as periphery of the wing disc. The method described here helps in quantifying concentration gradients in 3D, while keeping in consideration the shape of the tissue.

### Elucidating 3D organisation of cells within the curved epithelium of wing discs

Rearranging the image datasets from microscope reference to the sample surface coordinates has several advantages. In this section, we utilise this method to describe the challenging task of understanding the 3D organisation of cells within a curved pseudostratified epithelium. Although the apical arrangement of cells is widely considered to be a proxy for 3D organisation of cells, a detailed description of the cell shape along the apico-basal axis is necessary to appreciate the physical principles of cell packing. Pseudostratified epithelia are a special kind of epithelia wherein the nuclear packing density is very high. This forces the cells to have staggered nuclear distribution with respect to their neighbours (Kirkland *et al*., 2020; Hecht *et al*., 2022), necessitating morphometric quantifications of cell shape along the apico-basal axis. In the wing discs, we observe that the 3D organisation of cells is highly complex. Examples of reconstructed cells (Methods) of this pseudostratified epithelium reveal a plethora of shapes (Figure 6A). Using k-means clustering (Methods), these shapes can be clustered into three categories based on the cross-sectional area changes along the apico-basal axis: cells tapered towards the basal surface (Inverted Bowling Pin); cells bulged towards the basal surface (Bowling Pin); or cells bulged in the centre with tapered apical and basal surfaces (Rolling Pin) (Figure 6B). The apical area alone cannot describe the 3D shape as all three groups have similar apical areas (Figure 6B).

**Figure 6:**
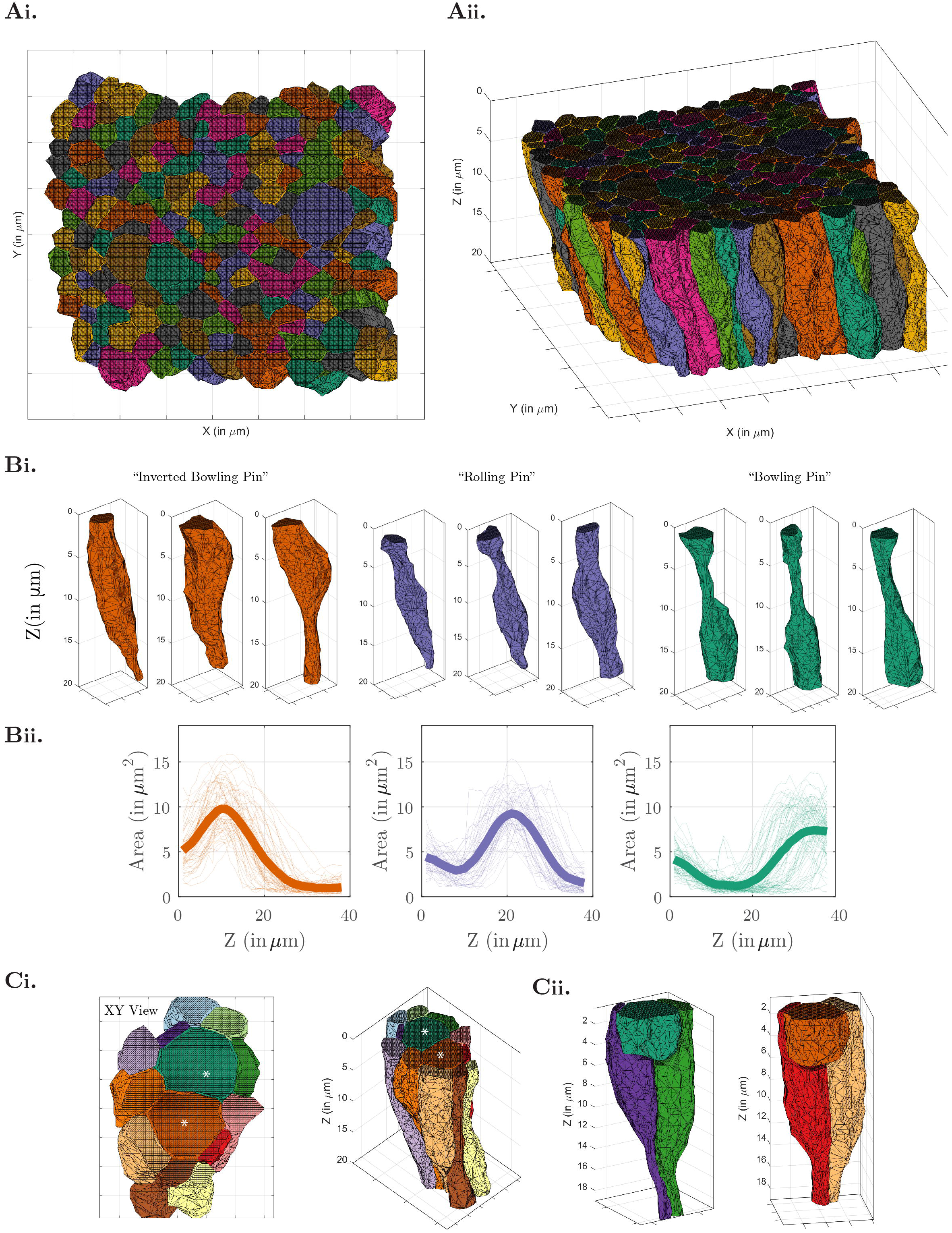
Neighbour exchanges observed along the apical-basal axis in 3D epithelium organisation. A: Cell boundary segmentation and cell surface mesh reconstruction along the apico-basal axis of cells shown as XY(i) and XYZ (ii) views. B: Pseudostratified cells show a diverse set of shapes. Based on cross-sectional area changes along the apico-basal axis, 188 cells (not including the dividing cells) were clustered into three categories using k-means clustering (i). While some cells show increased cross-sectional area at the top, some bulge in the centre and some towards the basal side. These were intuitively named as Inverted bowling pin (orange), Rolling pin (blue) and Bowling pin (green). The area changes across the apico-basal axis are indicated. Mean is shown as a thicker line (ii). C: Dividing cells show increased Apical area and are restricted to the 5-10 μm in depth on the apical side (i). The cells surrounding the dividing cells occupy the volume below the dividing cells (ii).

Other than the three categories of cells described above, a fourth category encompasses dividing cells. A conserved feature of metazoan cell division is change in cell shape, with cells rounding up before the mitotic spindle assembly and reverting to the resting state shape after cytokinesis (reviewed in (Cadart *et al*., 2014)). Similarly in the wing disc, dividing cells are found in the apical part of the epithelium, which has a larger cross-sectional area (Figure 6Ci) (Meyer, Ikmi and Gibson, 2011). The 3D view of dividing cells shows dramatic apical translocation, and these cells are observed within 5-6 μm depths from the apical surface. Additionally, the shapes of neighbouring cells are transformed to accommodate dividing cells by occupying the volume beneath (Figure 6Cii).

### Evaluating neighbour exchanges along the apico-basal axis

To further understand the details of cell organisation, we segmented 327 cells and measured cross-sectional area changes along the apico-basal axis (Methods). The cell arrangement of cells on the apical surfaces is more uniform, with a tight distribution of apical area (Figure 7Ai, 7Aii). The distribution of cross-sectional area becomes broader and bimodal along the basal surfaces (Figure 7Aii). On the basal side, cells have both larger and smaller cross-sectional areas compared to the apical areas, as expected from the observed cell shapes of the pseudostratified tissue. Within each basal surface, cells with smaller cross-sectional areas are constricted while those with larger areas form a bulge. The coefficient of variation of cross-sectional area increases from apical to basal surfaces.

**Figure 7:**
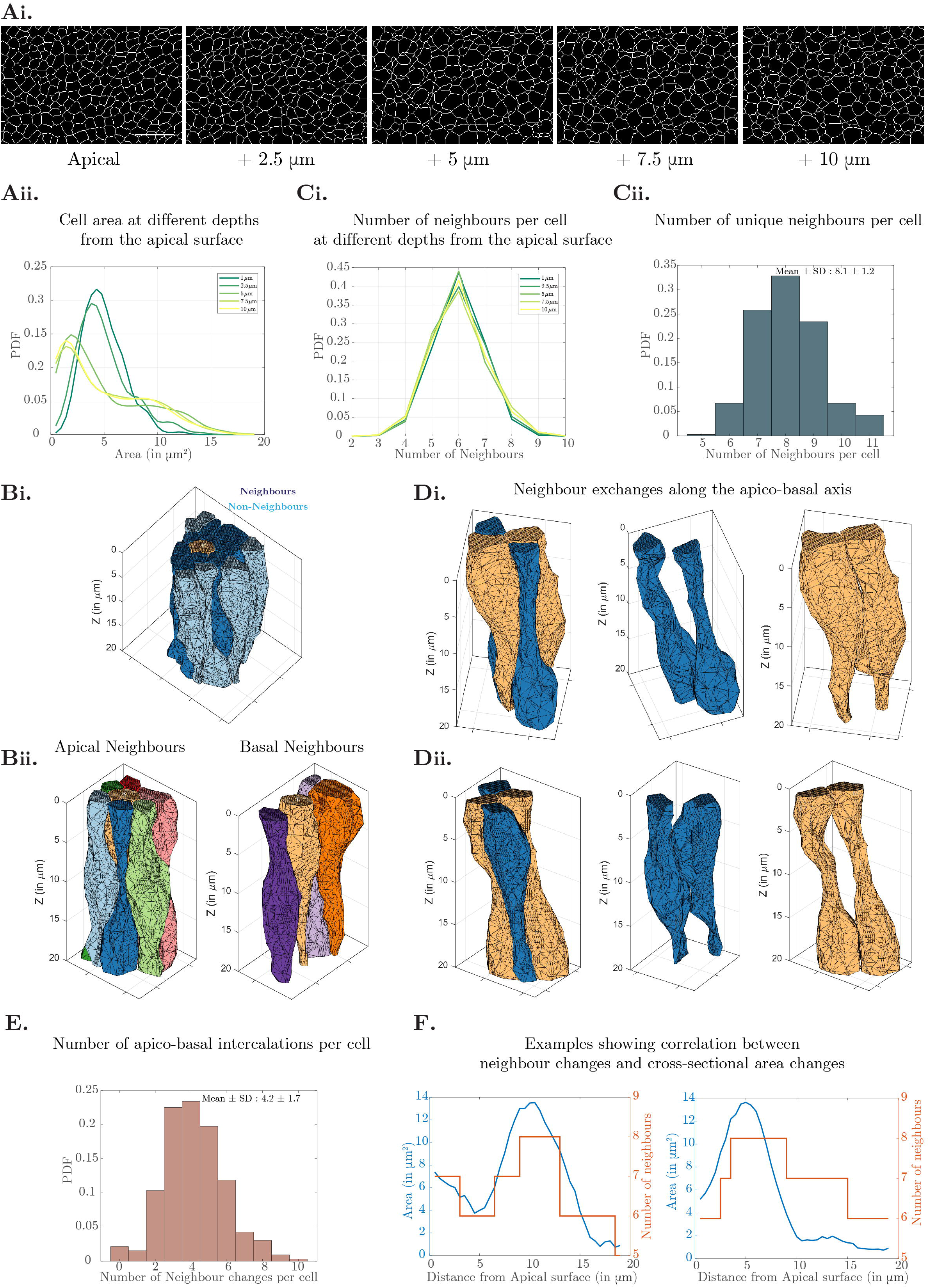
Neighbour exchanges observed along the apical-basal axis in 3D epithelium organisation. A: Segmented cells (i) and the probability density function of the cross-sectional area of cells (ii) is shown at different depths from the apical surface. Compared to the apical plane, at larger depths from the surface, two populations of cells are observed. While the apical cell area is a tighter distribution, distribution of cell area at 5μm from the apical surface is broader reflecting cell populations which are smaller than apical area and bigger than apical area. N=329 reconstructed cells from two samples. B: Cell neighbours are seen at distances of one cell diameter away on the apical side (i). These neighbours form contacts with the cell along the apico-basal length (ii). * indicates the cell of interest. C: The number of neighbours per cell at different depths from the apical surface averages at six neighbours per cell at all the depths (i). Probability density of unique neighbours per cell, including apical neighbours as well as neighbours observed along the apico-basal axis, averages at 8.1±1.2 neighbours. N=329 reconstructed cells from two samples. D: Different styles of the neighbour exchanges are observed. The blue cells depicted in both examples are not neighbours at the apical side but form neighbours in the basal planes (i) or in the medial planes (ii). E: Probability density function of number of neighbour changes/apico-basal intercalations observed per cell. On an average, each cell has 4.2±1.7 (Mean ± SD) intercalations. N=329 reconstructed cells from 2 samples. F: Area changes along the apico-basal length is positively correlated with the number of neighbour changes.

At all depths, the average number of neighbours per cell is six (Figure 7Bi). However, the frequency distribution of neighbour numbers on the apical and the basal sides are not identical, suggesting that the neighbour relationships could change along the apico-basal axis (Figure 7Bi). Indeed, on an average, cells have eight unique neighbours (Figure 7Bii, 7Ci). We even observed cells with 11 different neighbours along their apico-basal axis. As a result of having many neighbours, cells are in direct physical contact not only with the cells which are neighbours on the apical side but even with cells that appear to be two cell diameters away on the apical side (Figure 7Cii).

How are these cells then packed? We observe that some cells which are neighbours on the apical surface separate along the basal surface, allowing those which were not neighbours on the apical to intercalate along the medial or basal planes (Figure 7D). These neighbour exchanges, called pseudo-T1s or apico-basal intercalations, are frequently observed along the apico-basal axis. On an average, four such neighbour exchanges can be seen for each cell along the apico-basal axis (Figure 7E). The neighbour exchanges also positively correlate with the cross-sectional area changes. An increase in cross-sectional area results in an increase in the number of neighbours and vice versa (Figure 7F). The variability in cross-sectional areas along the apico-basal axis is a key determinant in defining the neighbours. The methods detailed here can be used to further explore the dynamics of cell shape changes within complex tissue architectures, during growth accompanied by cell divisions and cell death.

## Discussion

Studying cell fates and cellular organisation within curved tissues in 3D has long been a challenging problem. Quantitative analysis of fluorescently labelled curved tissues entails a comprehensive understanding of the interaction of light (imaging) with matter (tissue). This requires a systematic deconstruction of the tissue morphology into tailored geometric coordinates, considering the sample morphology, the imaging modality and reference imaging axes. Using the first principles-based methodology described here, the transformation of high-resolution confocal images planes from the frame of reference of the optical axis of the microscope into a laminar organisation with respect to the outermost sample surface, provides access to detailed quantitative analysis. Our data-based intensity correction procedure, which corrects the intensity variations brought about by sample geometry, provides a coherent method for evaluating the intensity distribution of biochemical signals in apical and basal layers. In addition, this type of transformation permits the detailing of cell shape and cell-packing within curved tissues. The morphometric measurements have important consequences for understanding the physical principles of cell packing as well as the generation of biochemical cues in the form of gradients or fields.

Defining the physical principles of cell packing is key to understanding the formation of curved tissue. Our careful approach accounting for tissue geometry has uncovered distinct cell shapes of cells within the pseudostratified epithelium of the *Drosophila* wing disc. Based on the area changes as a function of apico-basal axis, cells were clustered into three categories. These various cell shapes could be formed due to high nuclear densities. Interkinetic nuclear migration is a feature of pseudostratified cells (Strzyz, Matejcic and Norden, 2016). In vertebrate pseudostratified neuroepithelium, nuclear migration is initiated during G2 stage (Norden *et al*., 2009; Baffet, Hu and Vallee, 2015). In contrast, in the pseudostratified epithelium of *Drosophila*, the movement of nucleus to the apical surface is observed just prior to the M phase (Mandaravally Madhavan and Schneiderman, 1977; Gibson *et al*., 2006; Kirkland *et al*., 2020; Hecht *et al*., 2022). During other phases of the cell cycle, there is no concomitant match between the location of the nucleus and the cell cycle stage (Meyer, Ikmi and Gibson, 2011). As shown here, the dividing cells are apically translocated, while the surrounding neighbours occupy the volume beneath dividing cells. It would be interesting to evaluate how neighbours undergo shape changes during cell division events, while keeping the tissue intact along both apical and basal surfaces.

Given the challenges in visualising cells in 3D, most studies have focussed on understanding cell neighbour relationships using apical markers (Baena-López, Baonza and García-Bellido, 2005; Classen *et al*., 2005; Gibson *et al*., 2006; Aigouy *et al*., 2010). However, the apical arrangement need not be a proxy for cell organisation along the apico-basal extent of the tissue. In fact, we find that although predominantly hexagonal cells are found along all layers across the apico-basal axis in disc proper cells, the number of unique neighbours per cell is more than those present at the apical side. Neighbour exchanges through apico-basal intercalations have been observed previously in columnar cells (Rupprecht *et al*., 2017; Sun *et al*., 2017; Gómez-Gálvez *et al*., 2018). However, the frequency of such intercalations is much higher for pseudostratified cells as shown here for wing disc (~4 transitions per cell) and recently reported in mouse embryonic explants (~5 transitions per tube cell) (Gómez *et al*., 2021) compared to simple columnar cells. Correlation between apico-basal cross-sectional area changes and neighbour changes (as shown here and recently reported in cells of mouse explants (Gómez *et al*., 2021)) indicates that the driving force for cell shape is a key determinant in neighbour relations. Pseudostratified epithelia are often seen as organ precursors and are highly proliferative tissues (reviewed in (Norden, 2017)). Given the dynamic nature of the tissue, it would be exciting to next understand the temporal frequencies of apico-basal cell intercalations and how such interactions contribute towards cell-cell juxtracrine signalling.

Finally, the interpretation of concentration profiles of biochemical signals is key for understanding cell fate specifications. Morphogen gradients, which provide positional information during development, have been extensively studied (reviewed in (Tabata and Takei, 2004)). Considerations in image acquisition and image analysis are essential to quantitatively estimate morphogen concentrations at different positions. Our quantification now provides detailed estimation of concentration gradients within a curved tissue where cell co-ordinates along the apico-basal, anterior-posterior, and dorso-ventral axis are clearly classified. This allows estimating apical, medial, and basal concentration profiles across the dorso-ventral axis (considering *Wg* as a reference plane as considered here), as well as assessing the extent of variability in morphogen gradients (Iyer *et al*., 2022). The method can also be applied to estimate concentration profiles across the anterior-posterior axis by considering *Dpp* producing cells as a reference plane. As described here, fluorescence intensity variations due to scattering linked to sample geometry is maximally seen for outer layers of a curved epithelium. The intensity variations due to sample geometry should be deconvolved to determine biologically relevant concentration profiles. Apically localised morphogens will therefore be subject to such imaging artefacts and careful evaluation of the sample geometry with the imaging frame of reference is needed to extract the (near) accurate profiles of varying biochemical signals. The methods described here will undoubtedly serve to accurately delineate morphogen distributions, necessary for understanding the mechanisms behind the role of biochemical and mechanical cues in building 3D tissues.

## Acknowledgements

We thank JP Vincent (Crick Institute, UK) for sharing Wg-GFP flies. We thank the Central Imaging and Flow Cytometry Facility (CIFF) for imaging. We acknowledge Thomas Lecuit, Kabir Hussain, Subhash Sadhu, Sivaramakrishnan Swaminathan and members of Mayor laboratory for discussions related to this project. CP acknowledges NCBS-TIFR graduate fellowship and Company of Biologists’ Travelling Fellowship. SM acknowledges J.C. Bose Fellowship from SERB-DST, Government of India, and India Alliance DBT – Wellcome Trust Margdarshi fellowship (IA/M/15/1/502018). MR and SM acknowledge support from the Department of Atomic Energy (India), under project no. RTI4006. MR acknowledges support from the Simons Foundation (Grant No. 287975) and acknowledges the award of JC Bose Fellowship from SERB-DST, India. The work in the lab of TES was funded by a NRF Fellowship NRF2012NRF-NRFF001-094, core funding from the Mechanobiology Institute, National University of Singapore and start-up support from Warwick Medical School, University of Warwick.

## Author Contributions

CP, SM, TES conceived the study. CP conducted the experiments. CP and TES worked on the analysis pipeline. CP analysed the experimental data. KSI, MR conducted the theoretical analysis. CP, SM and TES wrote the manuscript with inputs from KSI and MR.

