## Supplementary Information for "Quantitative analysis of three-dimensional cell organisation and concentration profiles within curved epithelial tissues"

### Supplementary Note 1:

*Understanding the dependence of cell orientation on intensity:*

In curved tissues, cells are oriented approximately parallel to the local surface normal of the manifold describing the curved tissue surface. In such a scenario, the orientation angle of cells, $\theta$ - defined as the angle between the normal vector on the apical surface of cells and the vertical ($z$) illumination axis - contributes to a reduction in fluorescence intensity from the cell surface, and therefore the recorded concentration of the surface molecules. This curvature effect, if not accounted for in the image analysis, can lead to erroneous results. This can be simply illustrated as follows. Consider a curved tissue under observation. The apical surface of cells within this tissue, positioned at vertical distance $z_0$ from the objective plane, with orientation angle $\theta$, can be depicted as a square of side $a$ whose surface normal is tilted away from the $z$-axis by an angle equal to $\theta$ located at distance $z_0$ away from the objective (Figure S4A(i)). The intensity of the light emitted from the apical surface would incur a power law decay with the distance to the objective plane. For a given orientation angle $\theta$, this distance is $z_0 + a.\sin(\theta)$. The average intensity $\langle I \rangle$ from the apical surface of this region can then be calculated as

$$\begin{aligned} \langle I \rangle &= \frac{1}{a^2}\int_0^a dx \int_{z_0}^{z_0 + a\sin\theta} \frac{\alpha}{z^{n+1}}\frac{dz}{\sin\theta} \\ &= \frac{\alpha}{an\sin\theta}\left(\frac{1}{z_0^n} - \frac{1}{(z_0 + a\sin\theta)^n}\right) \\ &\simeq \frac{\alpha}{z_0^{n+1}}\left[1 - \frac{n+1}{2}\frac{a\sin\theta}{z_0}\right] \end{aligned} \tag{1}$$

Eq.1 and Figure S4Aii illustrate that the average intensity recorded from the apical surface of a region within the curved tissue depends on its orientation angle. The orientation effect is more pronounced if this region lies at a shorter distance from the recording plane. At larger distances, the contribution of reduction of intensity due to Z depth will be larger than the effect of cell orientation.

In the above explanation, we have implicitly assumed that the apical surface of cells within the tissue is fluorescently labelled and recorded. This need not be the case in general as the localisation of the fluorescently labelled biomolecule could be anywhere within the cell. Even in such a case, the trend with orientation angle will remain unchanged. However, what may affect this trend is if fluorescently labelled regions of interest are restricted to a thin stripe (like the apical surface, apical endosomes) or distributed across broad intracellular regions (like the nucleus). To ascertain this, we considered two extreme scenarios - one in which the molecules are localised to a thin narrow region within the cell (Figure S4B(i)), and another in which the molecules are distributed across the volume of the cell (Figure S4C(i)). For these cases, we included the details of microscopy and estimated the intensity recorded as $I(\vec{r}')$

$$I(x', y', z') = \int \frac{e^{-\frac{(x-x')^2}{\omega_x^2} - \frac{(y-y')^2}{2\omega_y^2} - \frac{(z-z')^2}{2\omega_z^2}}}{(z_0 - z)^2} c(\vec{r})d\vec{r} \tag{2}$$

where $\vec{r} = (x,y,z)$ and $c(\vec{r})$ is the function for distribution of the molecules of interest, which is taken as a constant (set equal to 1) between two parallel surfaces with orientation angle $\theta$ and angle with respect to the $x$-axis as $\varphi$. The two surfaces are described by the equations: $x \tan(\pi-\theta) \cos(\varphi)+y \tan(\pi-\theta) \sin(\varphi)+z =$

$c_1$ and $x\tan(\pi-\theta)\cos(\varphi)+y\tan(\pi-\theta)\sin(\varphi)+z = c_2$ with $c_2 = -c_1 = 0.025$ for the thin slab (Figure S4B(ii)) and $c_2 = -c_1 = 1$ for the case of thick slab (Figure S4C(ii)). In this calculation, the numerical constants associated with the anisotropic Gaussian kernel of the microscope were set to be: $\omega_x = \omega_y = \frac{\omega_z}{2.05} = 0.24$. The z thickness of the thin slab is less than $\omega_z$ and the same for the thick slab is greater than $\omega_z$. Figures S4B(iii), S4C(iii) show the dependence of sample orientation along with *z-depth* on intensity. The contribution of orientation matters in the thin slab but not in the example of a thick slab. In the case of a thick slab, the loss in intensity for any sample orientation is predominantly because of the *z-depth*. Meanwhile, in the case of a thin slab, both sample orientation and *z-depth* contribute to the recorded intensities.

### Methods:

*Fly stocks and imaging:*

Fly stocks used in this study are wildtype *w1118*, CAAX-GFP (Kyoto DGRC - 109823), Wg-GFP (Port *et al.*, 2014), Rab7 eYFP (BDSC 62545). Flies were reared on corn flour medium containing sugar, yeast and agar along with antibacterial and antifungal agents. Flies were grown in 25°C incubators with 12-hour light/dark cycles for experiments and otherwise maintained at 18°C or 22°C.

The following primary antibodies were used: anti-E-Cadherin (1:20, DCAD2, DSHB), anti-Wg (1:100, 4D4, DSHB) and anti-Arm (1:20, N27A1, DSHB). Anti-Wg antibody was purified from mouse hybridoma and labelled using NHS-ester chemistry (Alexa Fluor 568 NHS Ester, A200003, Invitrogen). Labelled antibody was separated from free dye using column chromatography. DAPI was used for nuclear labelling.

Third instar larvae (96-110 AEL) were washed twice with distilled water first and twice with phosphate buffer saline (PBS). Larvae were transferred to watch-glass containing Graces complete media (G9771, Sigma) for wing disc dissection (Dye *et al.*, 2017). Dissected discs were fixed using 4% PFA at RT for 20 minutes. Fixed discs were washed with 1X PBS and mounted in imaging chambers.

To label the production plane, either endogenously labelled Wg-GFP wing discs were used or for other genotypes, wing discs were pulsed with anti-Wg-AF568 for 30 mins or 1 hour. For endocytic pulse, dissected wing discs were labelled on ice for 40 minutes in graces complete media with Anti-Wg-AF-568. Following this, wing discs were pulsed with fresh media containing Anti-Wg-AF-568 at 25°C. After the defined time of pulse, wing discs were washed thrice with ice cold graces complete media and thrice with ice cold 1X PBS. Discs were fixed on ice for 5 minutes, followed by incubation at RT for 15 minutes. Fixed discs were washed with 1X PBS before mounting and imaging.

Immunostaining was done according to published protocols. Briefly, fixed wing discs were washed with 1X PBS and blocked with 1X PBS + 10mg/ml BSA for 30 minutes at RT. Primary antibodies were diluted in blocking buffer. Discs were incubated with primary antibody overnight at 4°C. Discs were then washed thrice with 1X PBS and incubated with blocking buffer containing corresponding fluorescently tagged secondary antibodies (1:200 dilution, Jackson Laboratories) for 1-2 hours at RT. Labelled discs were washed and mounted in imaging chambers.

Simple imaging chambers were made on coverslip bottom dishes using double-sided tape as spacers. Wing discs were mounted in between the spacers with the peripodial membrane oriented closest to the coverslip. A porous membrane (Whatman, 7060-2513) was adhered on top of the discs using the double-sided tape spacers to keep the discs in place. Excess 1X PBS buffer was added in the dish and imaged. We thank Natalie Dye and Suzan Eaton for sharing the composition of live imaging media as well as the idea of using a membrane to hold the discs in place. The usage of membrane along with double-sided spacer helped in imaging the disc in three-dimensions with minimal deformations.

Discs were imaged on FV3000 laser-scanning confocal microscope using 60X/1.42 NA oil objective with acquisition pixel dimensions of 0.138 µm or 0.208 µm and Z stacks of size 0.4 µm or 0.5 µm.

*Defining the wingless production plane, Q, Perpendicular plane, R, and reference surface, S, of the wing disc*

The hand drawn 3D mask marking the area of interest (*i.e.*, columnar cells in the wing pouch) was first smoothened by 3D gaussian filtering with a sigma of 2 pixels. Subsequently, the outlines of the mask were linearly interpolated to extract the apical-most surface, *S*. This point cloud *S* is a three-dimensional curve in space. The point cloud was further processed to identify equidistant points on the curved surface, using the function *interparc* (D'Errico, 2021b). This set of equidistant points (5000 for every wingdisc) describes the apicalmost surface, *S* of the wing disc.

Wg is produced by a stripe of cells at the dorso-ventral boundary of the wing imaginal disc. The production domain is approximated to be the best-fit plane, denoted by *Q*, passing through the points of high intensity in a few selected slices. A plane in three dimensions can be described by the following equation:

$$ax + by + cz + d = 0$$

with atleast one of the numbers, $a, b, c$ being non-zero. A plane can also be described by a point and a vector that is perpendicular to the plane. If $\overrightarrow{P_0} = (px, py, pz)$ is a point on the plane and $\vec{n} = (a, b, c)$ is the orthogonal vector, then the above equation can be rewritten as:

$$ax + by + cz + d = 0$$

$$d = -(ap_x + bp_y + cp_z)$$

$$d = -(\vec{n} \,.\, P_0)$$

Using the function *affine_fit* (Fangdi Sun, 2021), the production domain (plane *Q*) was determined.

The perpendicular plane, denoted by *R* was determined using *affine_fit* (Fangdi Sun, 2021) as the plane with orthonormal vector perpendicular to that of production plane *Q*.

*Estimating distances within the wing disc*

The 3D stack images were acquired with pixel dimensions of 0.138-0.207 $\mu m$ X 0.138-0.207 $\mu m$ X 0.4-0.5 $\mu m$ in X, Y and Z respectively. For evaluation, the image was divided into averaging voxels of 1.5-2.2 $\mu m$ X 1.5-2.2 $\mu m$ X 0.5 $\mu m$ dimensions. On an average, 200000-300000 evaluation voxels constitute each wing disc. Each voxel was characterised by its centroid, $P\ (x_p, y_p, z_p)$. For each voxel within the region of interest, three distances were calculated:

- The distance from the apical surface, $D_{PP'}$, is the distance of point, *P*, to the nearest point *P'* on the surface, *S*.
- The distance from the production domain, $D_{PQ}$, is the distance of point, *P* to the plane, *Q*.
- The distance from the perpendicular domain, $D_{PR}$, is the distance of point, *P* to the plane, *R*.

In addition to the distances, for each evaluation voxel, median intensity from all the pixels within the voxel was calculated for each channel.

The distance $D_{PP'}$ was computed using a function *distance2curve* (D'Errico, 2021a). The surface, *S*, was approximated to be a series of small line segments (piecewise linear chordal segments). Euclidean distance of point P to these piecewise line segments was calculated. The minimum of this set was estimated as the distance $D_{PP'}$, with *P'* being the nearest point on the curve.

The production plane, *Q*, passed through point $P_Q$ and with normal vector, $\vec{n}$. The dot product of vectors $\overrightarrow{PP_Q}$ and $\overrightarrow{n_Q}$ defined the shortest distance, $D_{PQ}$, of point *P* from plane *Q*. Similarly, the perpendicular plane, *R*, passed through point $P_R$ and with normal vector, $\overrightarrow{n_R}$. The dot product of vectors $\overrightarrow{PP_R}$ and $\overrightarrow{n_R}$ defined the shortest distance, $D_{PR}$, of point *P* from plane *R*.

For reconstruction of images according to layers, the distances computed for each evaluation voxel was interpolated to all voxels. This was done using functions *interp3* and *griddata*. The interpolated distances were binned into 0.5 µm thick layers. Each layer was projected onto a 2D plane for visualisation as seen in Movies S4 and S5 and Figure 2E, 3A.

*Estimating surface normal and included angle between surface normal and illumination axis*

The point cloud of points describing the outermost surface, *S* was used to estimate surface normals using MATLAB function, *pcnormals*. Based on the surface normal, the included angle between the surface normal and the illumination axis (Z axis) was calculated. The larger the angle, the more curved is the surface with respect to the illumination axis.

$$\theta = cos^{-1}\left(\frac{\vec{u} \cdot \vec{\iota}}{|\vec{u}| * |\vec{\iota}|}\right)$$

where, $\theta$ is the angle between the two vectors, $\vec{u}$ is the surface normal and $\vec{\iota}$ is the illumination axis vector and with bounds: $0 \leq \theta \leq 90°$.

For each evaluation voxel within the wing disc, the closest point on the surface point cloud was estimated and the corresponding surface normal along with the included angle was assigned.

*Quantifying 3D intensity profiles:*

Towards quantifying the intensity as a function of distance from producing cells, each disc was first divided into 5 layers using $D_{PP'}$ or *d*. Subsequently, each layer was further divided into 30 bins as a function of distance from producing cells ($D_{PQ}$ or *x*). The mean intensity within each bin was calculated as shown below:

$$I_{d,x} = \frac{1}{N_{d,x}} \sum_{voxels,v}^{v \in \Omega_d \,\&\, v \in \rho_x} I$$

$$NI_{d,x} = \frac{I_{d,x}}{I_{d=1,x=1}}$$

where,

$$\rho_x = \{voxels, v: D_{PQ,x} < D_{PQ,voxels} < D_{PQ,x+1}\}$$

$$\Omega_d = \{voxels, v: D_{PP',d} < D_{PP',voxels} < D_{PP',d+1}\}$$

- $\rho_x$ is a set of all voxels with distance from production plane ($D_{PQ}$) ranging between (x) and (x+1)
- $\Omega_d$ is a set of all voxels with distance from surface ($D_{PP'}$) ranging between (d) and (d+1)
- $N_{d,x}$ is total number of voxels, v, within the set $\Omega_d$ and set $\rho_x$
- $I_{d,x}$ is mean intensity of all voxels, v, within the set $\Omega_d$ and set $\rho_x$ and $NI_{d,x}$ is the normalised intensity

All intensities were normalised by the intensity of bin, *d*=1 and *x*=1 to facilitate comparison across multiple samples. Each layer (*d*) is colour coded as a range from green to yellow with green representing apical (closest to the surface) and yellow representing basal (farthest from surface). For each disc, the mean and SEM is represented as a thick line plot. Lighter coloured lines represent intensity within individual bins parallel to the R plane.

*Construction of Normalisation Matrix and correction of intensity:*

2D and 3D normalisation matrices were constructed using intensities from multiple samples. For each disc, intensity of CAAX-GFP was divided by the mean intensity to allow comparison across experiments. The intensities from voxels belonging to a specific *z-depth*, specific $\theta$ (included angle between surface normal and illumination axis) and specific layer were tabulated. The mean of intensities belonging to each $(d, Z, \theta)$ was used to construct the normalisation look-up-table.

For intensity correction, for all evaluation voxels within the wing disc, $(d, Z, \theta)$ was estimated and divided by the correction factor from the normalisation matrix corresponding to the voxel's $(d, Z, \theta)$.

$$I'_P = \frac{I_P}{N_{d,Z,\theta}}$$

where,

- $I_P$ and $I'_P$ is the raw and corrected intensity of the evaluation voxel
- $N_{d,Z,\theta}$ is the correction factor from the normalisation matrix corresponding to the specific $(d, Z, \theta)$

*Comparing the geometry of hemi-ellipsoid and wing disc with respect to illumination axis*

A hemi-ellipsoid with semi axis length of 100, 50 and 50 µm was simulated using *ellipsoid* function. The surface normal for the surface point cloud of the hemi-ellipsoid was estimated using *pcnormals*. Using the surface normal, the included angle ($\theta$) between the surface normal and the illumination axis (Z axis) was calculated. *z-depth* ($z$) is measured as the distance along the Z axis. An increase in $z$ showed a corresponding increase in $\theta$. For a hemi-ellipsoid, for every $z$, there is some distribution in $\theta$. However, for a sphere, a unique $\theta$ is associated with every $z$ ($z = r\cos\theta$) (where $r$ is the radius of the sphere). A similar relationship was computed for different wing discs. Due to the heterogeneity in geometry across minor and major axis in wing disc samples, here, for every $z$, there is larger distribution in $\theta$. Therefore, a 3D correction matrix using $d, z, \theta$ is required.

The effect of orientation on intensity is observed in lower layers, albeit to a lesser extent. This could be because scattering becomes a more dominant effect deeper into slices. Therefore, we find that correction using 3D normalisation matrix $(d, z, \theta)$ works best for those samples enriched in apical surfaces (like membrane signal, endosomal signal). Meanwhile for signals deeper in the tissue, like nuclear intensity, correction based on a 2D normalisation matrix $(d, z)$ or $(d, \theta)$ is sufficient.

*Segmentation, reconstruction of cells in 3D and cell shape classification:*

On arranging the 3D image stack into layers from the outermost surface (of $d$ = 0.5μm step size), the intensities were projected onto a 2D plane. Cell segmentation was done in 2D using Tissue Analyzer (Aigouy, Umetsu and Eaton, 2016) and manually corrected for scenarios when cells constricted along the apico-basal axis. The tracked output from Tissue Analyzer was exported into MATLAB for further processing. The outer surface of cells was reconstructed using *boundary* and *trisurf* functions in MATLAB. *Regionprops* function was used to estimate the area changes as a function of apico-basal axis. In each Z plane, for each cell, *delaunayTriangulation* and *voronoiDiagram* functions were used to classify the identities of neighbours and number of neighbours. *kmeans* clustering was used to classify the shapes of cells using the cross-sectional area across the apico-basal axis for each cell as input. The number of classes was defined as 4. Dividing cells, rolling pins, inverted bowling pins and bowling pins emerged as outcomes of kmeans clustering.

*Apical area, transverse area, and cell orientation estimation:*

For apical area estimation, 3D stacks of wing discs labelled with *E-Cadherin* was projected onto 2D plane (along Z axis) using StackFocuser (Umorin, 2006). For transverse area estimation, different transverse sections from wing discs labelled with CAAX-GFP were chosen for segmentation. Cells were segmented using Tissue Analyzer (Aigouy, Umetsu and Eaton, 2016) and the segmentation output was exported to MATLAB. *Regionprops* function was used to estimate the area, centroid, and long axis. Angle between the long axis of the segmented cells and the Z axis was computed.

To generate the depth map for maximum intensity of *E-Cadherin*, 3D stacks of wing discs labelled with *E-Cadherin* were smoothened using gaussian filtering, sigma = 2. The Z-position of maximum intensity for each (x,y) position was computed and represented as a heat map.

### Supplementary Figures and Legends:

#### Supplementary Figure 1

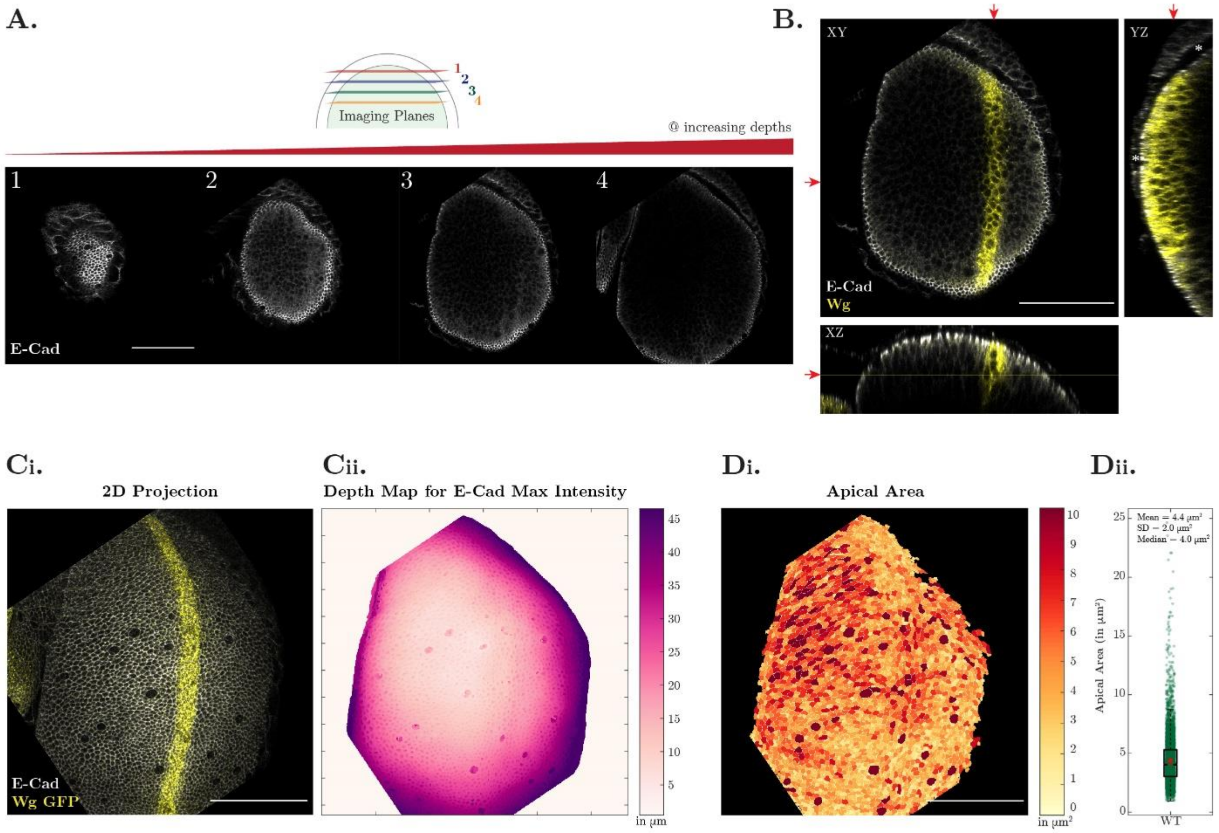

*Supplementary Figure 1: Apical Junction Marker, E-Cadherin, describes the outermost surface of cells embedded in the three-dimensions of the curved epithelium*

A: Z sections of *Wg-GFP* wing disc labelled with *E-Cadherin*. As the wing disc is dome shaped, each section has apical cells in the circumference of the image. The different Z sections show that each section has information about cells at different heights. For example, the rim of (4) has apical cells, while the central region depicts the basal part of the cell. B: Orthogonal projection of wing disc labelled with *E-Cadherin* and *Wg* of the Z slices shown in A. The red arrow marks the section represented in XY, YZ, XZ projection. The asterix highlights that the distance from peripodial cells and disc proper is not similar at different locations. C-D: 2D projection of wing disc labelled with *E-Cadherin* and *Wg* on XY plane (C) allows segmentation of cells and estimation of apical cell area (D, N = 2, n = 6009 cells). Depth Map shown in C(ii) denotes the Z position of maximum intensity of *E-Cadherin* for all cells.

### Supplementary Figure 2

**A.** **Bins using distances from Surface**

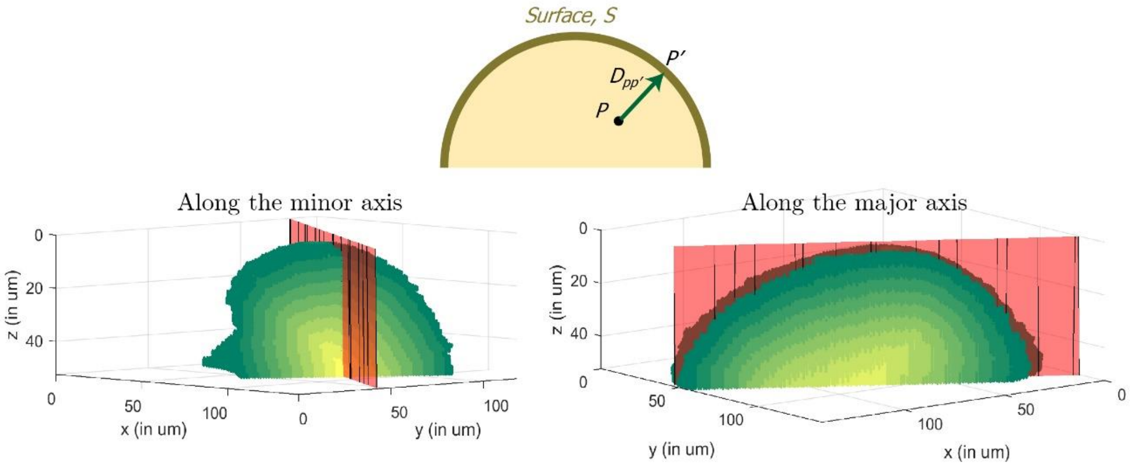


**B.** **Bins using distances from Production plane and Perpendicular plane**

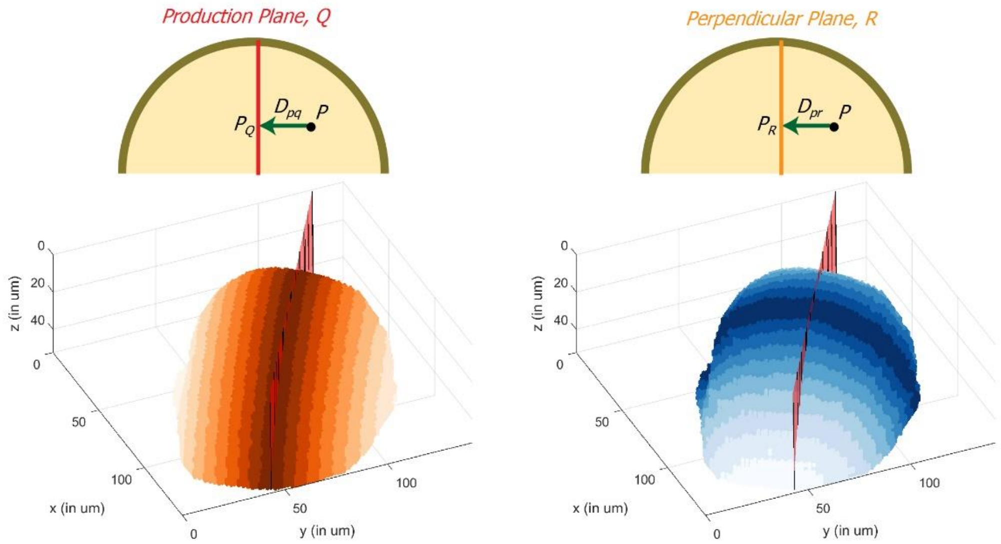

*Supplementary Figure 2: Binning wing disc based on distances*

A-C: Representative wing disc is binned based on computed distances: $D_{PP'}$ distance from the surface (A), $D_{PQ}$ distance from the production plane (B) and $D_{PR}$ distance from the perpendicular plane (C). Layers closest to the surface are marked in green and those farthest from the outer surface in yellow (A). Regions closest (and farthest) to the production plane are depicted in red (and white) (B). Regions closest (and furthest) to the perpendicular plane are marked in blue (and white) (C).

### Supplementary Figure 3

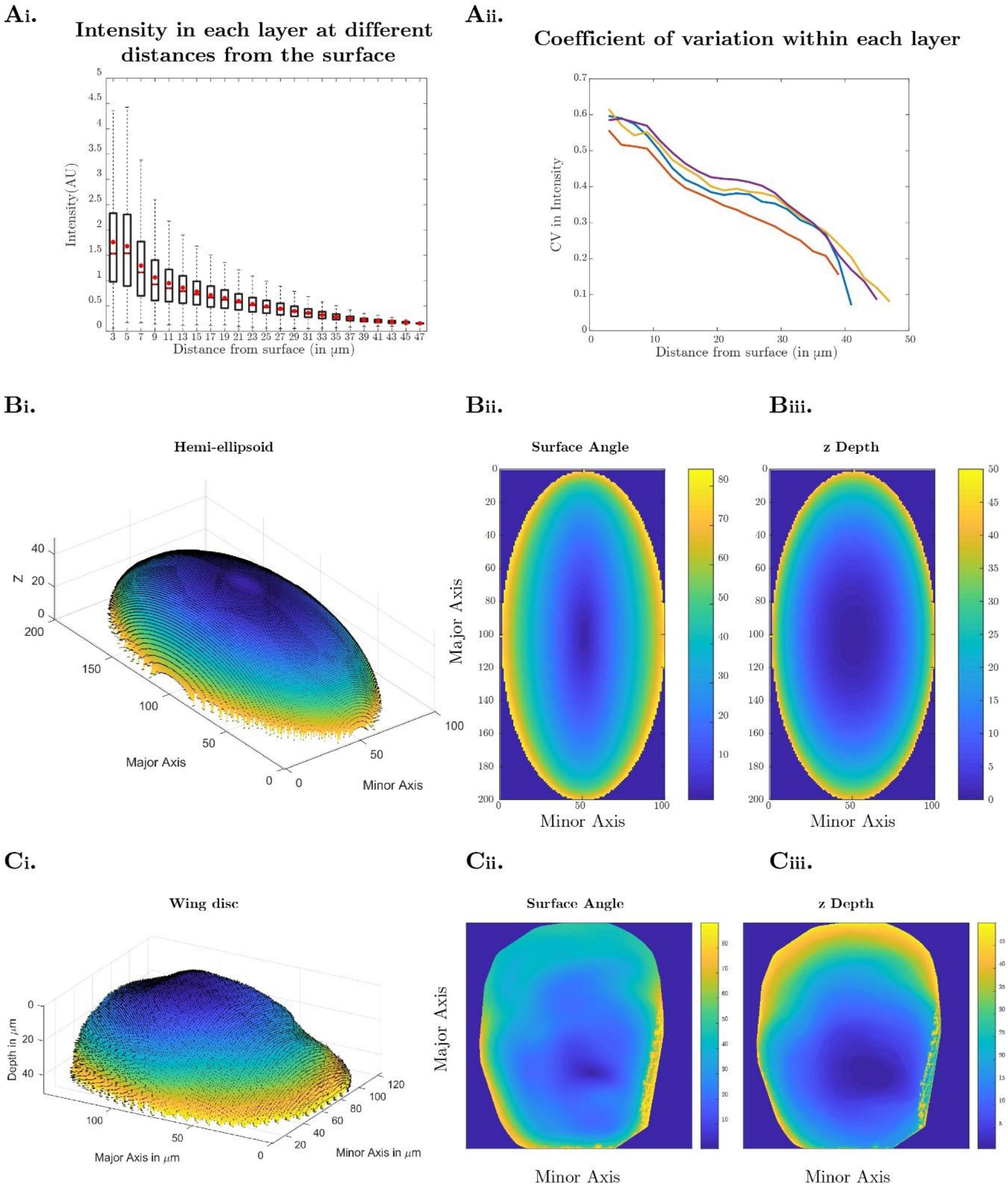

*Supplementary Figure 3: Effect of sample depth on intensity and comparing sample geometry parameters between hemi-ellipsoids and wing disc*

A: Normalised intensity (Intensity/Mean Intensity) of wing discs expressing CAAX-GFP shows that intensity reduces with increasing distance from the surface (i). The coefficient of variation in intensity shows that the variation is highest in the layers closer to the outermost surface (ii). The boxplot refers to intensity distributions from voxels identified in the topmost layer ($d$ = 1 is the apical most layer of 4.8µm thickness) of four wing discs. B-C: Surface of hemi-ellipsoid (B) and wing disc (C) with surface normals (i). 2D representation of surface angle (ii) and z depth (iii) for hemi-ellipsoid and wing disc show that the centre has lower surface angle, low z depth and the periphery has larger z depths, larger deviations in surface angles. The wing disc geometry is not as uniform as the hemi-ellipsoid. Both z depth and surface angle are required to describe the sample geometry of the wing disc.

**Supplementary Figure 4**

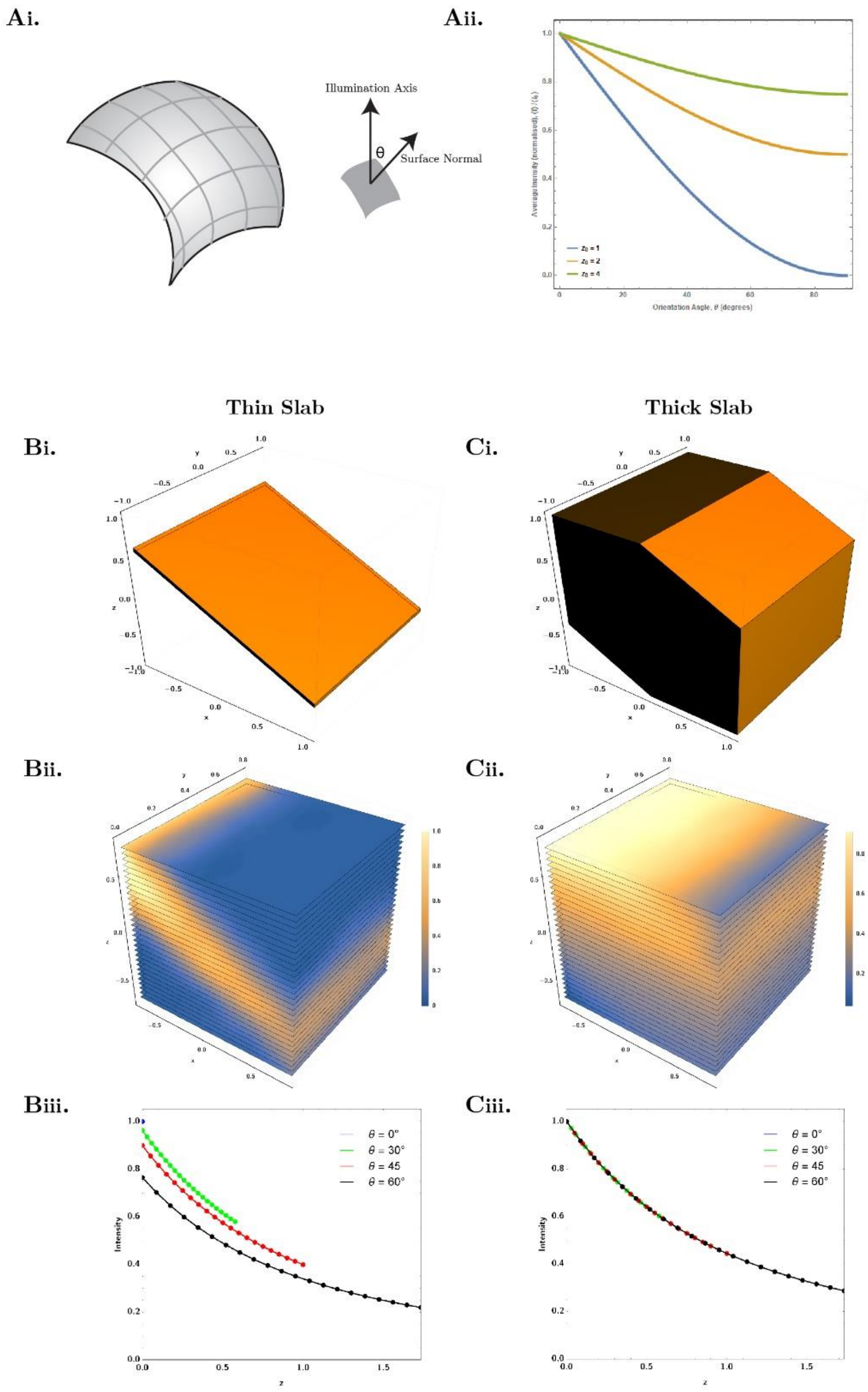

*Supplementary Figure 4: Understanding the dependence of cell orientation on recorded fluorescence intensities*

A(i): Apical surface of a cell (or a few cells) at a certain position along the curved tissue oriented at an angle $\theta$ with respect to the illumination (Z) axis. The average intensity recorded from the cell is computed as an integral of the decaying intensity over this surface. A(ii): The estimated intensity (see Eq.1) as a function of the orientation angle. The reduction in intensity is less pronounced when the cells are further away from the recording plane. B(i) and C(i): Narrow and broader distribution of fluorescently labelled markers within a cell considered as a constant function between two surfaces (see Supplementary Note 1). The orientation of these surfaces depict the local orientation of the cell. B(ii) and C(ii): Recorded intensity as per Eq.2 for the scenarios detailed in B(i) and C(i) respectively. B(iii) and C(iii): Recorded intensity from the points on the centre plane of the volumes described in B(i) and C(i) respectively, as a function of their z-distances from the recording plane. The distance of the origin from the recording plane in these figures is kept at $z_0 = 2$ (see Eq.2).

### Supplementary Figure 5

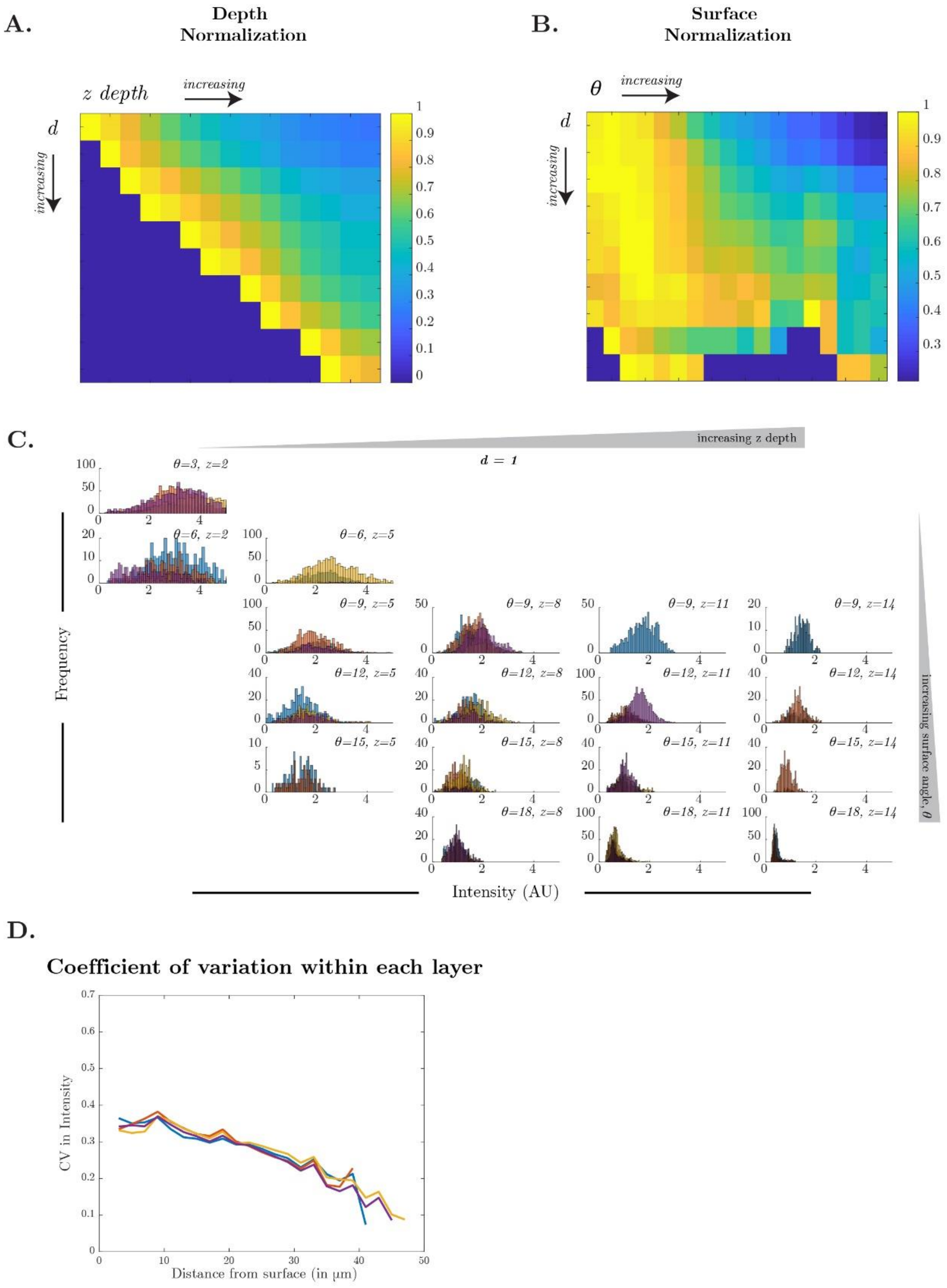

*Supplementary Figure 5: Normalisation matrices to correct the intensity bias brought about by sample geometry*

A-B: Depth-based normalisation matrix using ($d$, $z$) (A) or Curvature-based normalisation using ($d$, $\theta$) (B) as parameters were constructed using intensities from four wing discs labelled with CAAX-GFP The mean of intensities belonging to each ($d$, $z$) or ($d$, $\theta$) was used to construct the normalisation look-up-table.

C: An example of CAAX-GFP intensity distributions across wing discs (different colours in the histogram) for the apical layer ($d = 1$) at different ($z$, $\theta$). Intensity reduces with increasing $z$ and/or $\theta$.

D: Correction reduces the coefficient of variation of intensities observed within each layer. Compare this result with S3A(ii).

**Supplementary Figure 6**

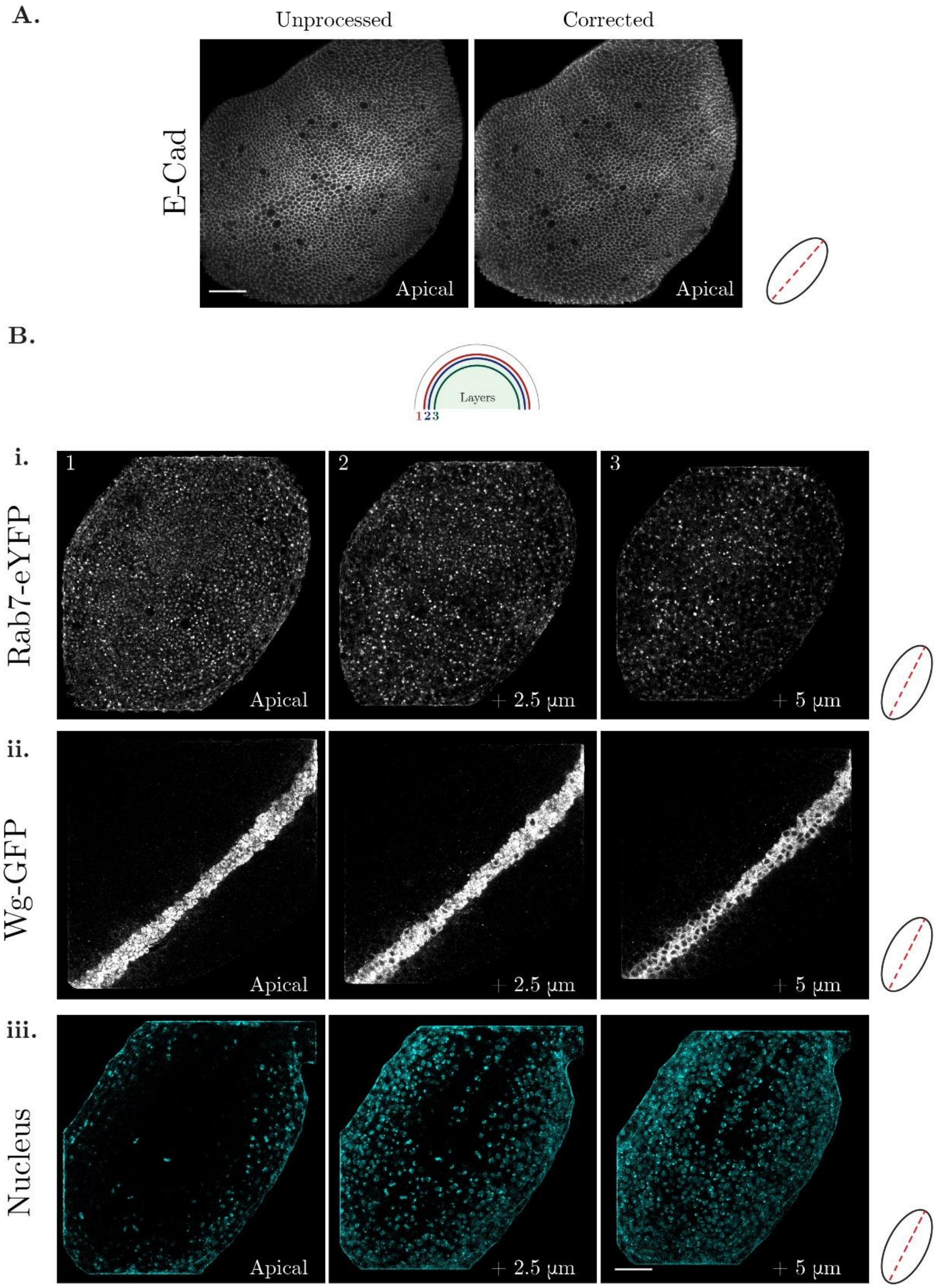

*Supplementary Figure 6: Some examples of intensity corrected layers*

A: Unprocessed and intensity corrected layers of wing disc labelled with *E-Cadherin*. B: Examples of intensity corrected layers of *Rab7-eYFP* (i), *Wg-GFP* (ii) and Nucleus (iii). Red line in the orientation schematic on the right shows the *Wg* producing cells. Scale bar: 20µm.

### Movie Legends:

Movie S1-S3: Movies with three different orientations of a representative wing disc expressing CAAX-GFP along Z axis (S1), Y axis (S2), X axis (S3). The axes and plane description are as shown in Figure 1. Scale bar: 10µm.

Movie S4-S5: Movies of raw (S4) and intensity corrected (S5) layers of wing disc expressing CAAX-GFP with each layer projected onto 2D plane. Each plane represents intensity information from a 0.5µm thick layer. Scale bar: 10µm (S4) and 20µm (S5). In S5, the intensity in each layer is normalised between 0 and 1.

Movie S6: Movie of wing disc labelled with apical marker (Armadillo in grey) and nuclear marker (DAPI in cyan) along Z axis. Scale bar: 20µm.
